# ESTROGEN IMPACTS NOD2-DEPENDENT REGULATION OF INTESTINAL HOMEOSTASIS

**DOI:** 10.1101/2025.07.04.663219

**Authors:** Mckenna Eklund, Edan A. Foley

## Abstract

Mutations in the innate immune receptor NOD2 are the greatest single genetic risk factor for Crohn’s disease, yet the mechanisms by which NOD2 regulates intestinal homeostasis remain unclear. We used a CRISPR-generated zebrafish model to determine the impacts of NOD2 deficiency on intestinal health. In cellular, molecular, and transcriptomic studies, we uncovered substantial effects of NOD2 deficiency on epithelial and immune compartments, including deregulated expression of developmental pathways that establish and maintain the gut epithelium, and an unexpected increase in the expression of multiple estrogen-response genes. In functional assays, we uncovered a mechanistic link between estrogenic signals and NOD2-deficiency phenotypes, whereby exposure to estrogen alone replicated the effects of NOD2-deficiency, and treatment with the estrogen receptor modulator tamoxifen reverted the epithelial defects observed in *nod2* mutants. Our findings identify a NOD2-estrogen regulatory axis that supports intestinal homeostasis and suggest that hormonal signaling may contribute to sex-specific aspects of Crohn’s disease.

## INTRODUCTION

Crohn’s disease (CD) is a chronic inflammatory disorder of the gastrointestinal tract with significant public health implications. The prevalence and incidence of CD continue to rise in North America and Europe, with particularly notable increases observed in developed and urbanized regions [1]. Mutations in the Nucleotide-binding Oligomerization Domain containing 2 (NOD2) locus are the most significant genetic risk factor for CD [2–5], increasing disease susceptibility by 15-40-fold and contributing to heightened risk of colorectal cancer [6,7].

Early studies proposed that NOD2 functions in Paneth cells to regulate antimicrobial peptide production, particularly α-defensins [8–11], in response to muramyl dipeptide (MDP) derived from bacterial peptidoglycan [12,13]. However, follow-up investigations suggested that defensin loss may be a consequence, rather than a cause, of CD, prompting a broader re-evaluation of NOD2’s intestinal function [14–16]. Emerging evidence indicates that NOD2 plays a role in regulating intestinal stem cell (ISC) dynamics, particularly under stress conditions [17]. ISCs express high levels of NOD2 [18], NOD2 is essential for stem cell proliferation in mouse models of T cell-induced enteropathy [19], and NOD2 is essential for microbe-induced epithelial regeneration in malnourished models [20]. Despite these insights, it is unclear if NOD2 supports ISC function and epithelial renewal under steady-state conditions.

To investigate how NOD2 regulates intestinal homeostasis, we used zebrafish (*Danio rerio*), a well-established and genetically tractable model of human intestinal biology and disease [21,22]. Zebrafish and mammalian intestines share conserved immune and developmental pathways [23–29], including key components of NOD2 signaling [30]. Importantly, zebrafish offer unique advantages for examining epithelial interactions *in vivo*: their optical transparency enables live imaging of intestinal architecture. Additionally, zebrafish replicate pharmacologically actionable CD phenotypes [31, 32].

In this study, we combined single-cell RNA sequencing, targeted gene expression analysis, and *in vivo* functional assays to investigate the involvement of NOD2 in intestinal epithelial proliferation, differentiation, and cell survival under homeostatic conditions. Our data uncovered a NOD2-estrogen axis that is critical for intestinal homeostasis and host responses to colitogenic stress, providing insights into the cellular mechanisms underlying NOD2-associated intestinal dysfunction and sex-specific variations in disease.

## RESULTS

### NOD2 deficiency reduces intestinal length but does not alter microbiota composition in zebrafish

To investigate the role of NOD2 in intestinal homeostasis, we used CRISPR-Cas9 mutagenesis to generate *nod2* mutant zebrafish that harbor a 4-base pair deletion in the first coding exon, resulting in a frameshift and premature stop codon that produces a severely truncated, non-functional NOD2 protein (Fig 1A). Adult *nod2^-/-^* zebrafish were homozygous viable and showed modest drops in body weight and standard body length by six months (Fig 1B-C). In contrast, *nod2^-/-^* adults exhibited an approximately 20% reduction in total intestinal length (Fig 1D). This reduction remained significant after normalization to either body length or body weight, indicating a genotype-specific decrease in relative gut size rather than a secondary effect of overall body scaling (Fig 1E-F). Histological analysis suggested architectural differences in the mutant intestine, including reduced epithelial fold height and altered mucosal organization (Fig S1), features that are examined in more detail in later figures. Using a custom-made rabbit anti-zebrafish NOD2 antibody, we confirmed intestinal NOD2 protein expression in 4-month-old adult WT intestines and validated the absence of functional NOD2 protein in *nod2^-/-^*mutants (Fig S1).

**Fig 1.**
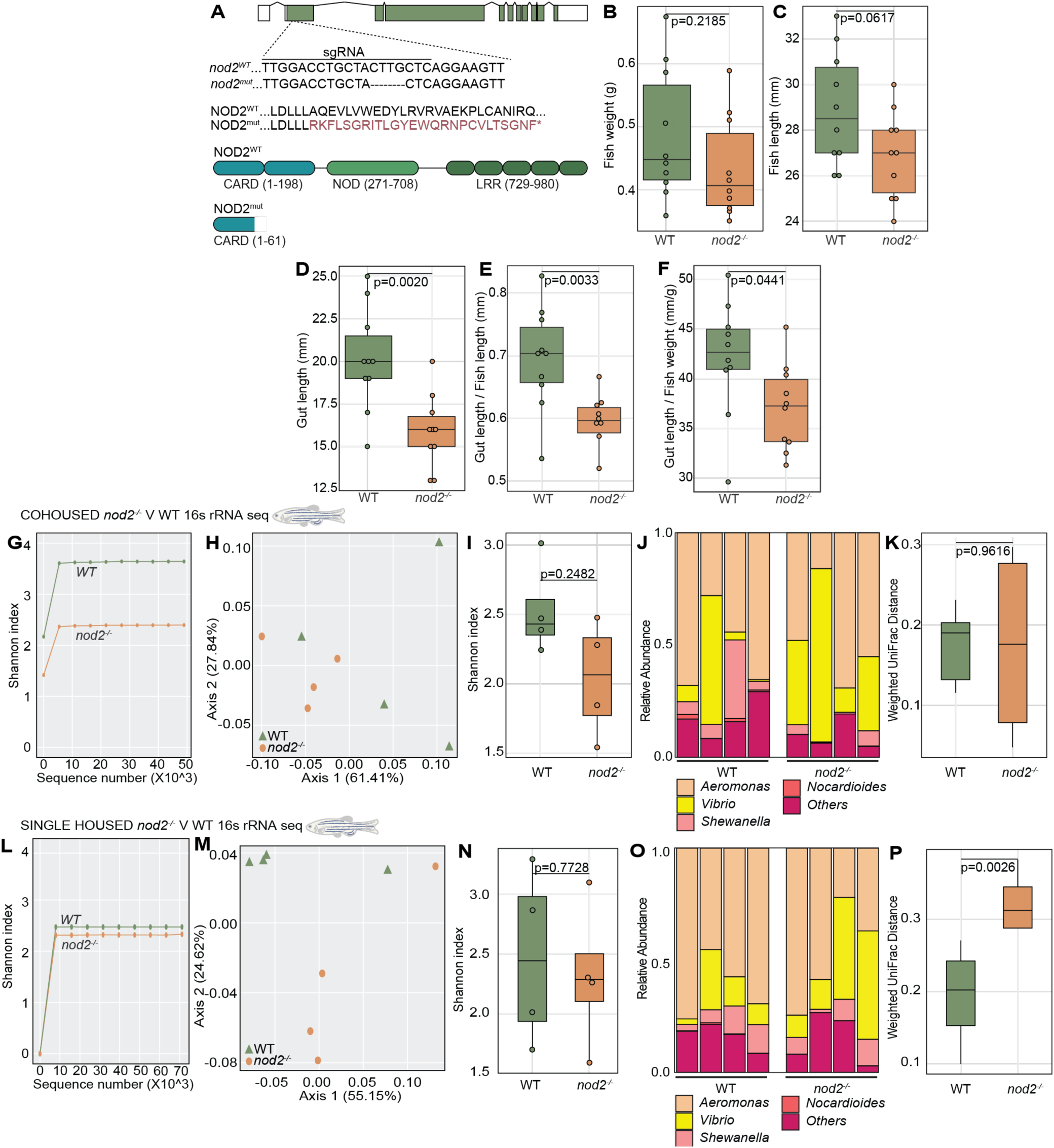
Generation of a nod2-deficient zebrafish model reveals reduced gut length without major microbiome disruption. **A)** Schematic illustration of the *nod2* transcript structure with all introns and exons (coding DNA in green). A sgRNA targeting exon 2 of nod2, within the region encoding the first CARD domain, was used to generate a 4-base pair deletion via CRISPR-Cas9. Measurements of **B)** fish weight, **C)** fish length, and **D)** gut length of 6-month-old adult WT and *nod2^-/-^* zebrafish in-crossed siblings; *nod2^-/-^*zebrafish exhibit significantly shorter gut lengths compared to WT, while body weight and length is comparable. **E-F)** Gut length normalized to body length **(E)** and body weight **(F)** showing reduced relative gut size in *nod2^-/-^* fish after size adjustment. **G-K)** Microbiota analysis of 6-month-old adult cohoused WT and *nod2^-/-^* intestines by 16S rRNA sequencing. Rarefaction curves (G), PCoA plot of beta diversity (H), Shannon index score of each genotype as a measure of alpha-diversity (I), relative abundance of dominant bacterial genera (J), and weighted UniFrac distances of beta-diversity (K) show no major differences between genotypes. **L-P)** Microbiota analysis of 6-month-old adult single-housed WT and *nod2^-/-^* intestines. Rarefaction curves (L), PCoA plot (M), Shannon index (N), relative abundance of bacterial genera (O), and weighted UniFrac distances (P) demonstrate comparable alpha diversity but genotype-associated differences in beta diversity and community composition. P-values were calculated using the Mann-Whitney U test. Each bar in relative abundance plots represents an individual intestine. Icon indicates the developmental stage at which the experiment was performed.

**Supplementary Fig S1.**
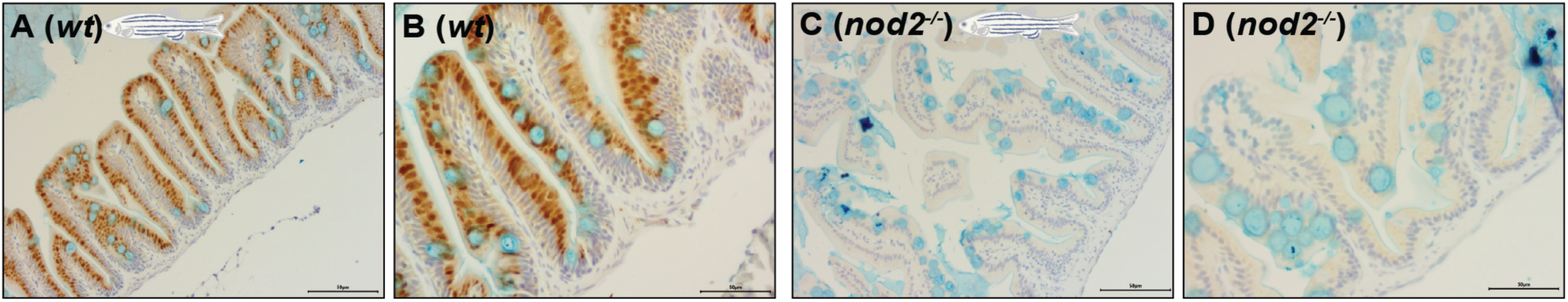
NOD2 protein expression in adult WT and *nod2^-/-^* intestines. A-D) Representative sagittal sections of 4-month-old cohoused WT and *nod2^-/-^* adult intestines immunostained with a custom anti-zebrafish NOD2 antibody. Images were taken at 20x (A, C) and 40x (B, D) magnification. n = 5 fish per genotype (mixed sex). Scale bars = 50 μm. Icon indicates the developmental stage at which the experiment was performed.

As interactions between NOD2 and sex are minimally defined, we initially used mixed-sex in-crossed sibling cohorts to define genotype-dependent intestinal phenotypes. Where later experiments identified sex-dependent variation, follow-up analyses were performed with sex recorded and analyzed explicitly. To determine if NOD2 deficiency modifies microbiome composition in zebrafish, we performed a deep-sequencing analysis of the V4 region of the bacterial 16S rRNA gene from 6-month-old adult WT and *nod2^-/-^* siblings. Under cohousing conditions, NOD2 loss did not have discernible effects on microbiota composition, representation, or diversity (Fig 1G-K). In contrast, when genotypes were maintained in separate tanks on the same recirculating water system under otherwise identical conditions, mutants showed comparable alpha diversity, but modest differences in beta diversity and relative taxonomic abundance relative to wildtype counterparts (Fig. 1L-P). These findings mirror data from mice [15, 33–35], and highlight the utility of fish for examining homeostatic roles of NOD2.

### High-resolution profiling of NOD2-deficient intestines uncovers early immune and epithelial alterations

To better determine the impacts of NOD2 deficiency on gut homeostasis, we performed single-cell RNA sequencing on whole intestines dissected from 6-month-old co-housed, WT and *nod2^-/-^* siblings derived from a heterozygous in-cross (2 males and 3 females in each group), effectively minimizing artefactual impacts of sex, environment and genetic variability. As zebrafish intestines are comparatively small but contain most cell types found in mammalian guts [23,27], this approach allowed us to characterize the consequences of NOD2 deficiency for all major intestinal cell types. In total, we generated high-quality transcriptomic data for 11,490 cells, that resolved into 13 distinct transcriptional clusters (Fig 2A, B) based on established markers from both zebrafish [23] and mammalian gut studies [27,36], including critical immune-regulatory epithelial, stromal, and hematopoietic lineages. Although *nod2* mRNA levels were below detection thresholds, our data revealed broad expression of NOD2 pathway components and signatures of activation across epithelial and immune subsets, closely paralleling expression patterns reported in the human intestine (Fig S2). Gene ontology analysis uncovered widespread transcriptional changes in *nod2^-/-^* intestines, with impacts on processes related to growth, development, and survival across epithelial and immune compartments, as well as modest shifts in cell type proportions, including apparent decreases in of goblet cell numbers and increases in leukocyte abundance (Fig 2C, D).

**Fig 2.**
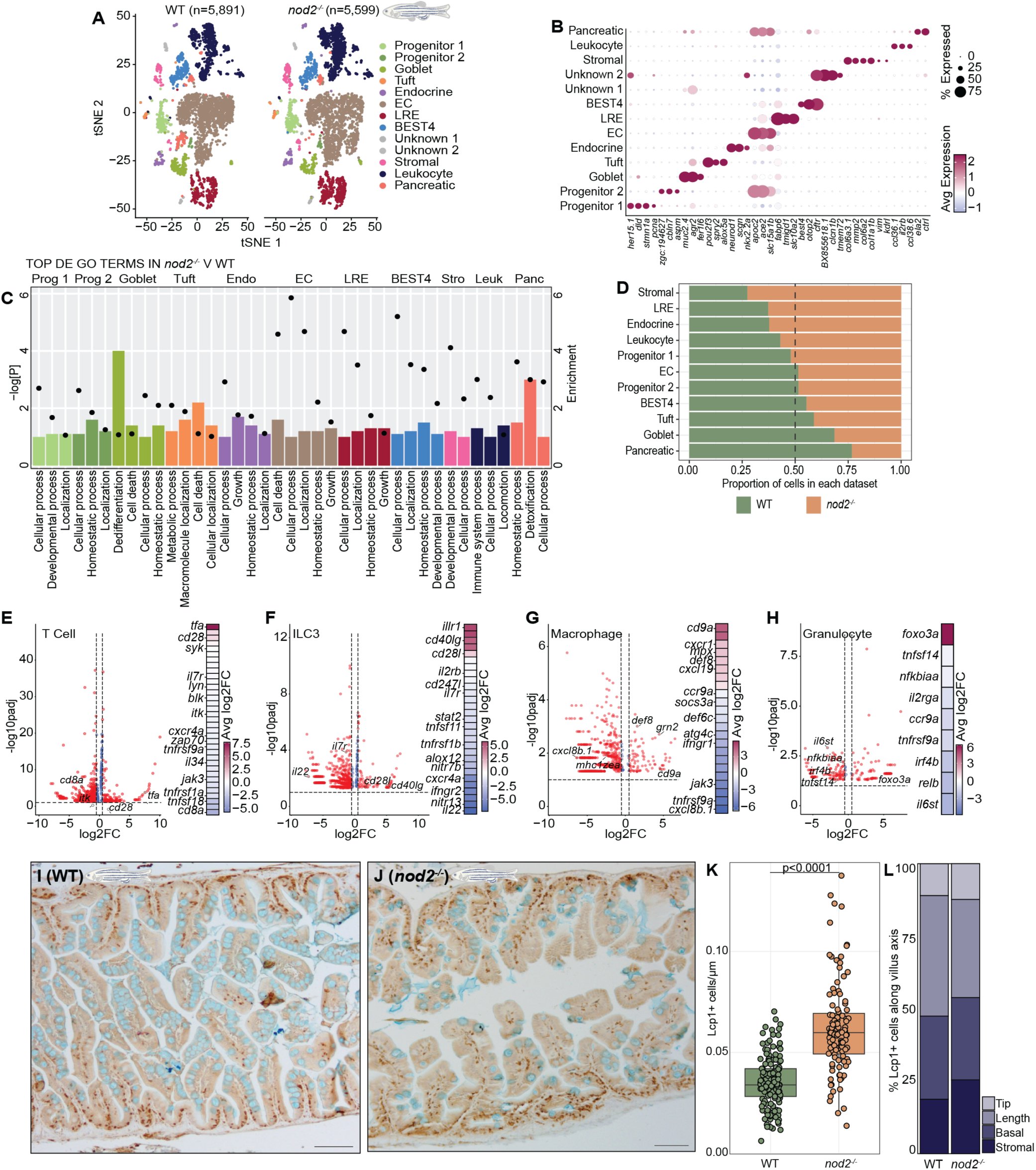
NOD2 deficiency alters intestinal epithelial and immune cell transcriptional programs. **A)** Individual 2D t-distributed stochastic neighbor embedding (tSNE) projections of 11,490 intestinal cells color coded by cell type, including progenitors, absorptive enterocytes (ECs), lysosome-rich enterocytes (LREs), goblet cells, tuft cells, stromal cells, and others. **B)** Dotplot representation of the expression levels of established markers in the indicated cell types. **C)** Representation of the top gene ontology terms associated with DE genes in *nod2^-/-^*intestines relative to WT sibling controls for each cell type. Relative enrichment scores are indicated by bar length and negative log P-values are shown with closed circles. **D)** Bar plot displaying the proportional representation of each cell type in WT and *nod2^-/-^*datasets. **E-H)** Volcano plots of DE genes in *nod2^-/-^* intestines relative to WT sibling controls in the indicated cell type (genes upregulated in *nod2^-/-^*appear on the right; downregulated genes appear on the left). Only genes with absolute log₂ fold change greater than 0.25 and adjusted p-value ≤ 0.05 are shown. Heatmaps display scaled expression values of top immune-related genes across each cell type. **I-J)** Immunohistochemical staining and **K)** quantification of Lymphocyte Cytosolic Protein 1 (Lcp1)^+^ cells per μm in co-housed 18-month-old WT and *nod2^-/-^* intestines. n = 5 fish per group; each point represents an individual measurement obtained from multiple images per fish intestine. Scale bars = 100μm. **L)** Spatial distribution of Lcp1⁺ cells along the intestinal fold axis, expressed as the percentage of total Lcp1⁺ cells localized to villus tip, villus length (mid), basal, and stromal compartments for each genotype. P-values were calculated using Mann-Whitney U test. Icon indicates the developmental stage at which the experiment was performed.

**Supplementary Fig 2.**
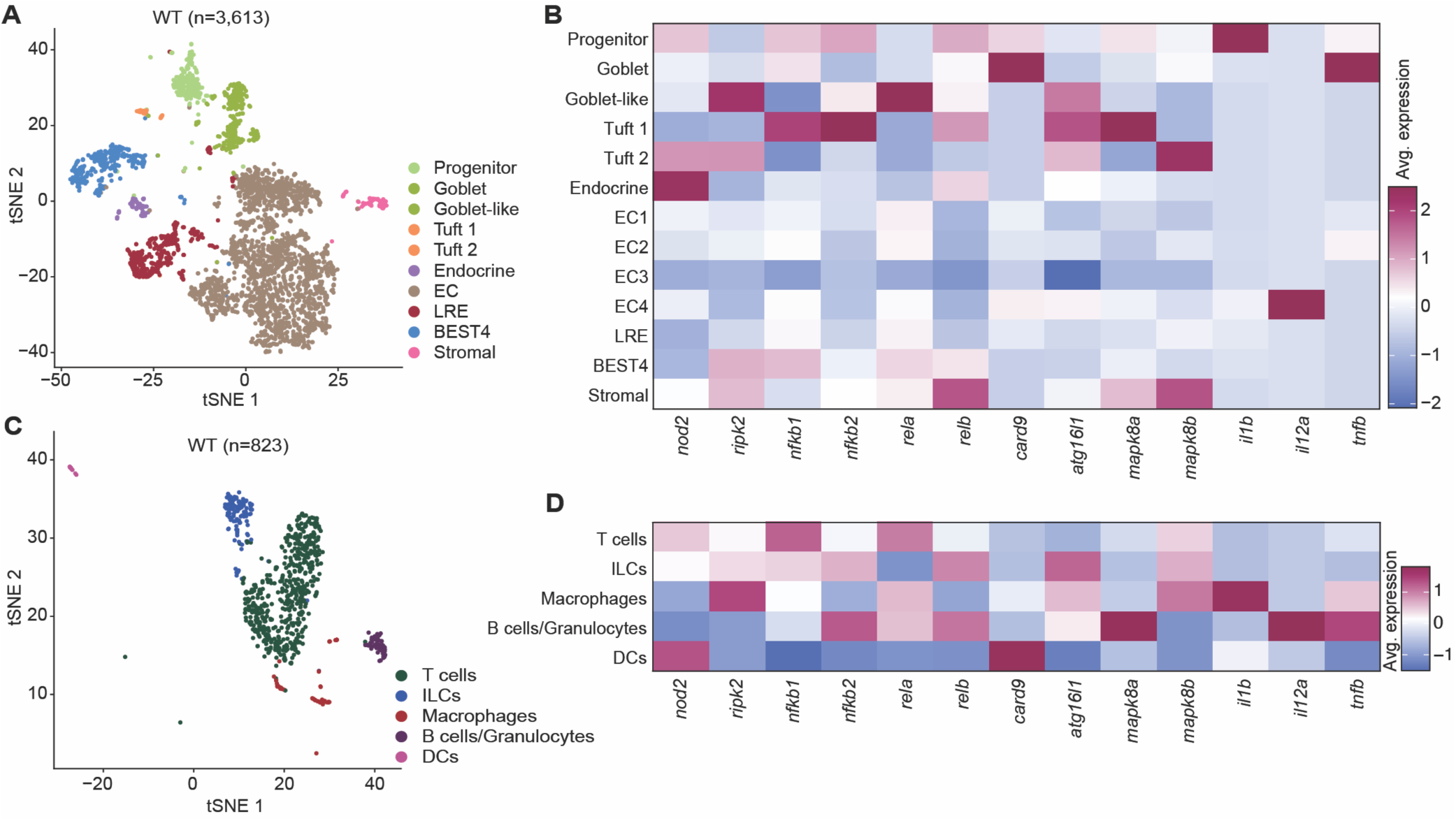
Single-cell expression of *nod2* across epithelial, stromal, and immune lineages in WT adults. **A-B)** tSNE plot (left) of single-cell RNA-seq from WT adult epithelial and stromal cells (n = 3,613), annotated by cell type. Heatmap (right) shows average expression of *nod2* and NOD2-pathway genes across epithelial and stromal clusters. **C-D)** tSNE plot (left) of single-cell RNA-seq from WT adult intestinal immune cells (n = 823), annotated by lineage. Heatmap (right) shows average expression of *nod2* and pathway-associated genes across immune subsets.

**Supplementary Fig 3.**
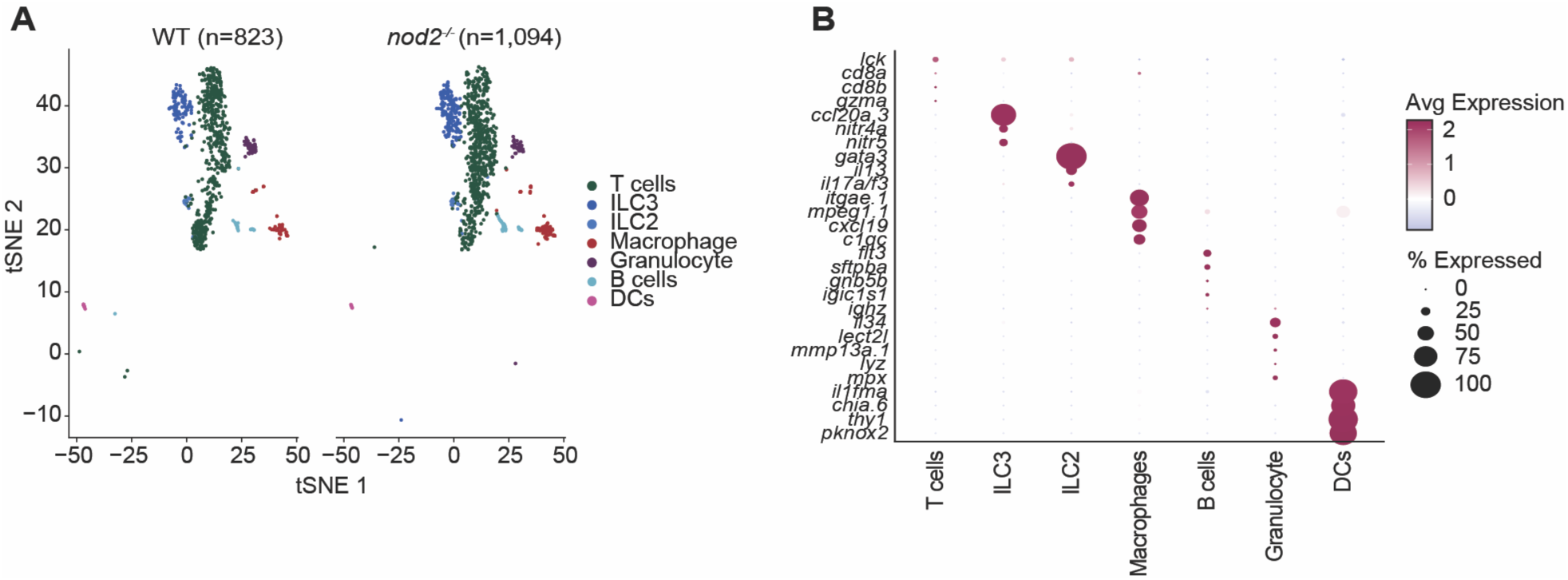
Identification of intestinal immune cell populations in WT and *nod2^-/-^*intestines. **A)** tSNE visualization of leukocytes from WT (left) and *nod2^-/-^* (right) zebrafish intestines (6-month-old, cohoused), colored by immune cell type based on transcriptional clustering and marker gene expression. Major immune subsets identified include T cells, ILC3s, ILC2s, macrophages, granulocytes, B cells, and dendritic cells (DCs). **B)** Dotplot representation of the expression levels of established markers in the indicated cell types from integrated data set.

**Supplementary Fig 4.**
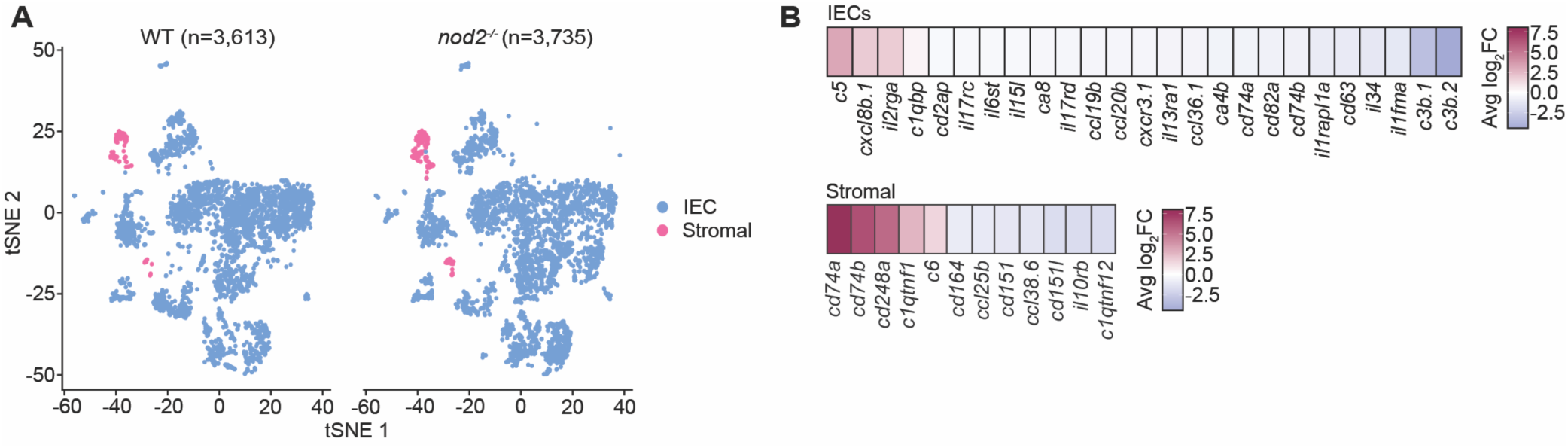
NOD2 deficiency enhances leukocyte-recruitment gene signatures in epithelial and stromal compartments. **A)** tSNE plots of intestinal epithelial cells (IECs; blue) and stromal cells (pink) from 6-month-old, cohoused WT and *nod2^-/-^*zebrafish. **B)** Heatmaps show differential expression of immune cell recruitment-associated genes in IECs and stromal cells (For all genes adjusted P-value ≤ 0.05).

Given the central role of gut-associated leukocytes in CD [37,38] and the high expression of NOD2 in intestinal immune cells [39–43], we elected to characterize effects of NOD2 deficiency on leukocyte transcriptional states in greater detail (Fig 2E-H). Gut-associated leukocytes were identified based on transcriptional clustering and canonical immune marker gene expression (Fig S3). We discovered notable impacts of NOD2 deficiency on expression of co-stimulatory and cytotoxic markers in T cells (Fig 2E); reduced expression of *il22* and *il7r* in ILC3s (Fig 2F); and dysregulated NF-κB signaling in granulocytes (Fig 2H). In macrophages, expression of the CD-linked cytokine *cxcl8b.1* was significantly downregulated, while fibrosis and autophagy associated genes (*cd9a*, *atg4c*) were upregulated (Fig 2G). Additionally, *nod2^-/-^* IECs exhibited altered expression of chemokines (e.g., *cxcl8a.6, ccl20b, cxcr3.1*) and cytokine signaling components (e.g., *il2rga*, *il17rc*, *il34, Fig S4A-B*), while stromal cells altered transcripts related to leukocyte activation and adhesion (e.g., *cd74a*, *cd248a, Fig S4A-B*), pointing to substantial, intestine-wide shifts in the immune landscape of NOD2-deficient adults.

To evaluate the *in vivo* consequences of these transcriptional shifts, we used immunohistochemistry to quantify the abundance of cells that express the pan-leukocyte Lcp1 marker in 18-month-old NOD2-deficient guts and WT controls (Fig 2I, J). Consistent with the widespread impacts of NOD2 deficiency on leukocyte transcriptional states, we observed a significant accumulation of Lcp1⁺ cells in aged *nod2^-/-^* intestines relative to wildtype controls (Fig 2K), confirming that NOD2-deficienct intestines harbor a larger population of functionally dysregulated immune cells.

To define the effects of NOD2 deficiency on leukocyte accumulation within the intestine, we stratified Lcp1⁺ cells by anatomical position along the intestinal fold axis, separating villus tip, villus length (mid), basal, and stromal compartments (Fig 2L). This spatial analysis showed that the increase in Lcp1⁺ cells in *nod2^-/-^* intestines was distributed across epithelial and stromal regions rather than confined to a single niche, consistent with broad immune compartment remodeling rather than a purely localized infiltrate.

### NOD2 regulates intestinal epithelial progenitor dynamics

Alongside anticipated effects on immune-regulatory leukocytes, we noted dysregulated gene expression in NOD2-deficienct epithelial lineages, including prominent effects on pathways linked to cell growth, development and survival (Fig 2C). Given the role of the intestinal epithelium in preserving immune homeostasis, we asked if NOD2 modifies epithelial renewal. To answer this question, we focused on cellular features associated with tissue replenishment, starting with progenitor cell proliferation. Using our single-cell RNA sequencing data, we found that NOD2 deficiency significantly attenuated the expression of key growth regulators in the progenitor cell populations (Fig 3A), including critical elements of the Wnt/β-catenin (*wnt7ba*), TGFβ (*tgif1*), and growth (*myca*) pathways (Fig 3B), suggesting that NOD2-deficiency disrupts the molecular framework required for intestinal epithelial growth and renewal. Feature plots confirmed expression of the respective genes within progenitor populations, supporting their relevance to early epithelial renewal programs (Fig 3C, D).

**Fig 3.**
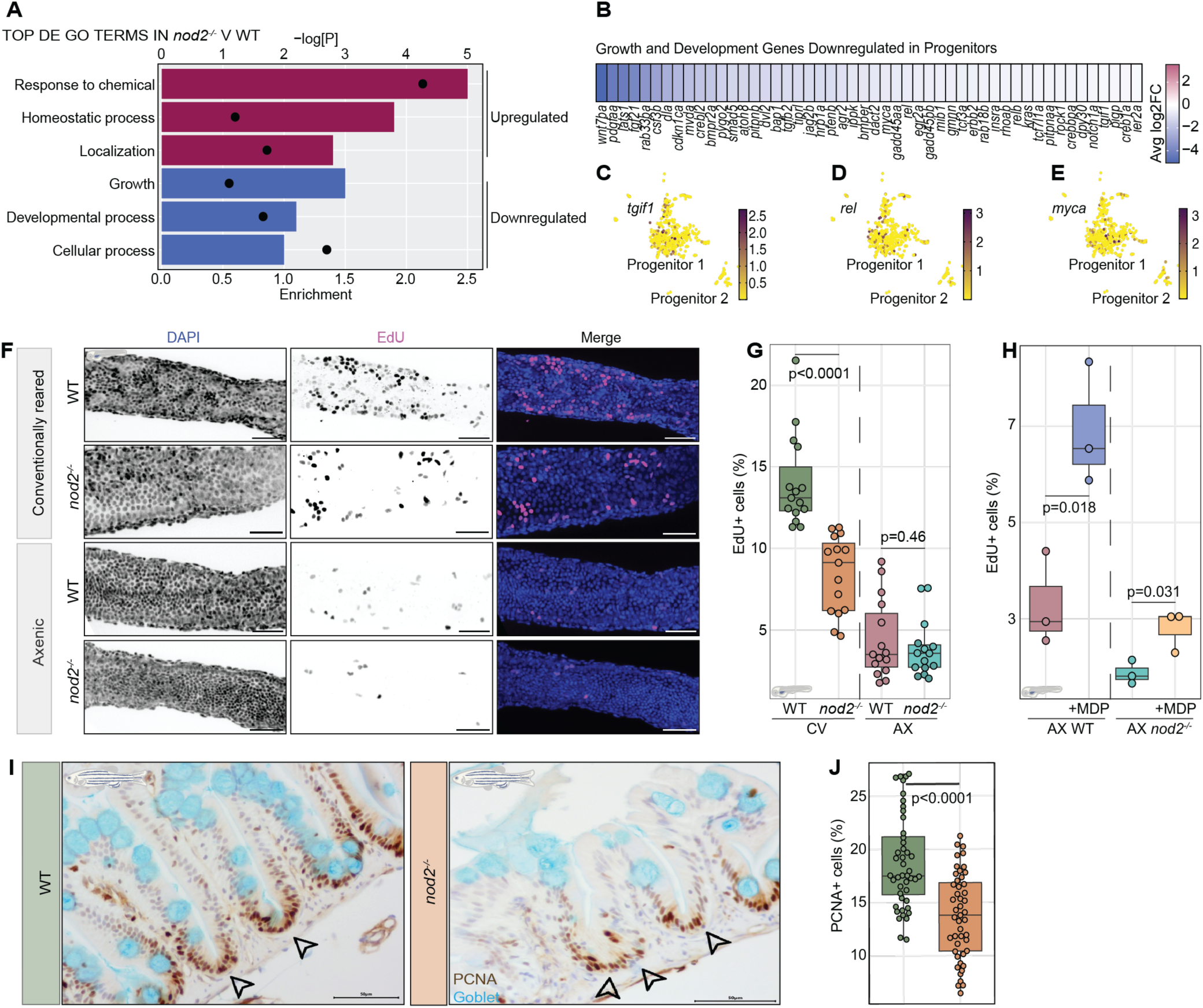
*nod2*-deficiency impairs intestinal growth and proliferation. **A)** The top three up, or downregulated gene ontology terms in *nod2^-/-^* progenitor cells relative to WT controls. Relative enrichment score for each term is indicated by bar length and negative log P-values are shown with closed circles. **B)** Heatmap of representative growth and development gene expression in *nod2^-/-^*progenitor cells relative to WT controls (for each gene adjusted p-value ≤ 0.05). **C-E)** Feature plots confirming expression of *tgif1* (C), *rel* (D), and *myca* (E) in progenitor cells. **F)** Immunofluorescence images of conventionally reared or germ-free WT or *nod2^-/-^* larval intestines (7 dpf), shown in rostral-to-caudal orientation from the mid intestine. In merged false-colored images, DNA is labeled in blue and EdU in magenta. Scale bars = 35μm. **G)** Quantification of the percentage of EdU^+^ cells in conventionally reared or germ-free WT or *nod2^-/-^*larval intestines (7 dpf), determined from whole-mount confocal images by counting EdU⁺ nuclei within a standardized intestinal region of interest. Each dot represents a measurement from a single fish intestine. **H)** Flow cytometry quantification of the percentage of EdU^+^ cells in germ free WT or *nod2^-/-^* larval intestines (7 dpf) following mock or MDP treatment (1 μg/ml), using dissociated intestinal cells. One dot represents 15 zebrafish larvae guts. **I)** Immunohistochemical images of a sagittal posterior intestine section from WT and *nod2^-/-^* intestines (6-month-old, cohoused adults) stained for Proliferating Cell Nuclear Antigen (PCNA-arrowheads). Sections were counterstained with Alcian blue (Goblet) and quarter-strength Hematoxylin Gill III in lavender blush (nuclei). Images are shown in rostral-to-caudal orientation. Scale bars are indicated in each panel. **J)** Quantification of % PCNA^+^ cells from WT and *nod2^-/-^* intestines (6-month-old, cohoused adults). Each dot represents a single villus measurement from n=5 zebrafish intestines per genotype. Scale bars = 100μm. P-values were calculated using Mann-Whitney U test or Kruskal-Wallis test. Icons indicate the developmental stage at which the experiment was performed (larval or adult).

**Supplementary Fig 5.**
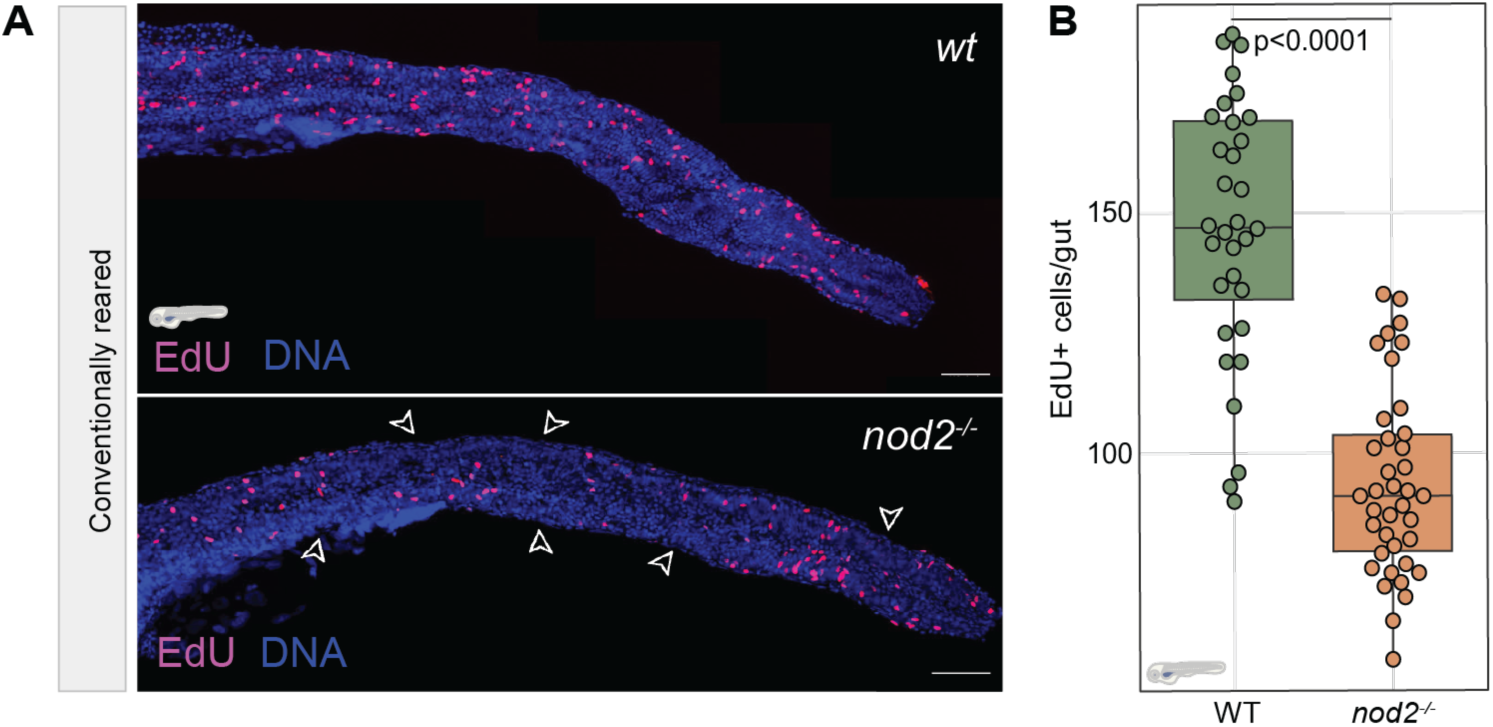
NOD2 is required for epithelial proliferation in the developing zebrafish gut. **A)** Immunofluorescence images of whole-mount intestines from conventionally reared WT and *nod2^-/-^*larvae (7 dpf), shown in rostral-to-caudal orientation from the base of the intestinal bulb to the cloaca. EdU⁺ cells (magenta) and nuclei (blue, DNA) are shown in merged false-colored images. White arrowheads highlight regions where *nod2^-/-^* intestines lacking EdU⁺ cells. Scale bars= 150μm. **B)** Quantification of total EdU⁺ cells per gut in WT and *nod2^-/-^* larvae (7 dpf), determined from whole-mount confocal images by counting EdU⁺ nuclei across the entire intestinal epithelium. Each point represents an individual larva; *nod2^-/-^*intestines exhibit a significant reduction in proliferating cells. P-values were calculated using Mann-Whitney U test. Icon indicates the developmental stage at which the experiment was performed.

**Supplementary Fig 6.**
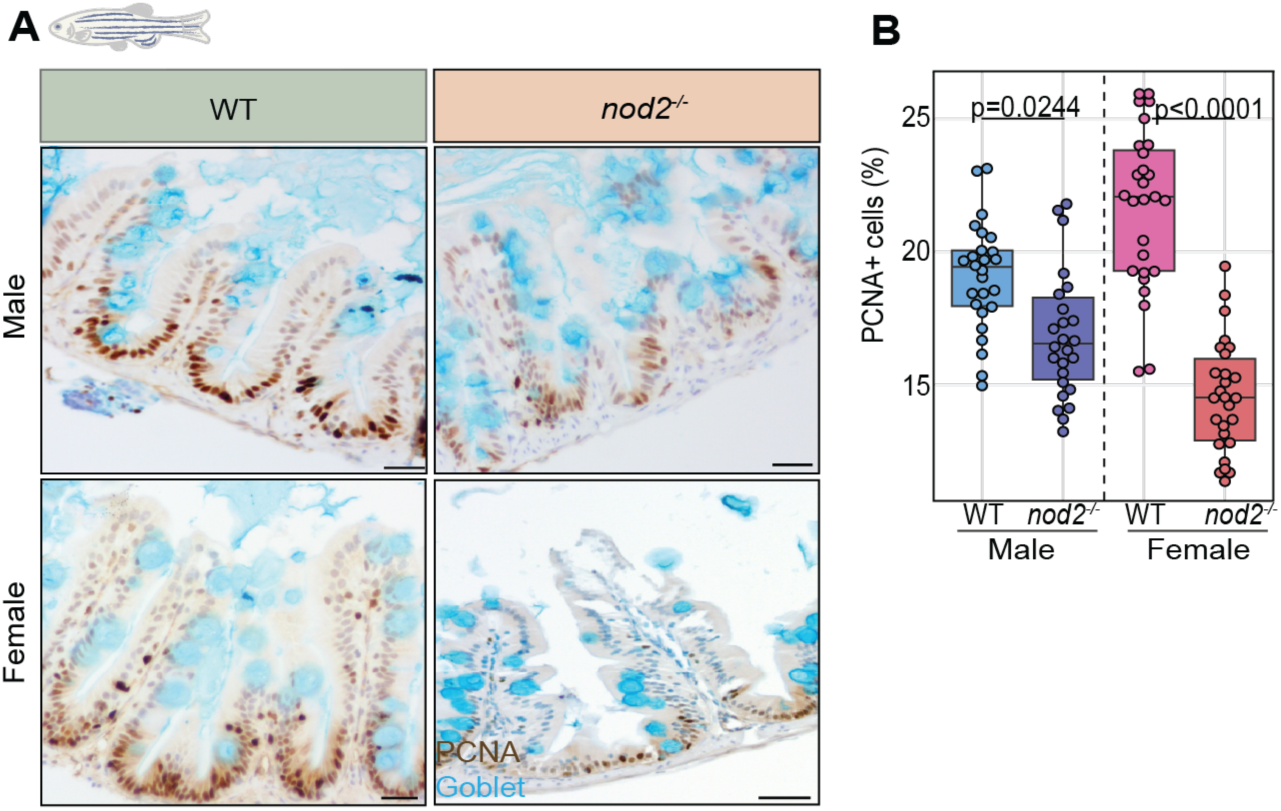
Loss of NOD2 decreases intestinal epithelial proliferation with sex-specific effects. **A)** Representative sagittal sections of posterior intestine from 6-month-old WT and *nod2^-/-^*zebrafish, separated by sex. Sections were stained for proliferating cell nuclear antigen (PCNA; brown) and counterstained with Alcian blue to visualize goblet cells (cyan). Images are shown in rostral-to-caudal orientation and were collected from comparable posterior intestinal regions using identical imaging settings. Scale bars = 20 μm. **B)** Quantification of epithelial proliferation shown as the percentage of PCNA⁺ epithelial cells per villus in WT and *nod2^-/-^* intestines, analyzed separately in males and females. Each dot represents a single villus measurement from n=5 zebrafish intestines per genotype. P-values were calculated using Mann-Whitney U test or Kruskal-Wallis test. Icon indicates the developmental stage at which the experiment was performed.

To test putative roles for NOD2 in epithelial cell proliferation, we quantified incorporation of the S phase marker EdU in intestines of WT and *nod2^-/-^*larvae at seven days post-fertilization. In this baseline comparison, we quantified EdU⁺ cells from whole-mount confocal images by counting EdU⁺ nuclei across the entire intestine, to obtain an unbiased whole-organ measure of proliferation. We discovered that NOD2 deficiency led to a 50% drop in the number of cycling cells (Fig S5A, B), confirming that NOD2 is required for epithelial proliferation *in vivo*.

To determine if this effect depends on microbial cues, we repeated the EdU labeling assay with seven-day post-fertilization axenic larvae alongside a conventionally reared control population. For these comparisons, we quantified EdU⁺ cells from whole-mount intestines by confocal imaging and counting EdU⁺ nuclei within a standardized intestinal region of interest (ROI), preserving tissue architecture and ensuring consistent quantification across rearing conditions. In agreement with a previous study [29], we found that wildtype larvae exhibited reduced proliferation under axenic conditions (Fig 3F, G). Furthermore, and consistent with data in Fig. S5, we observed a near 50% decline of EdU incorporation in conventionally reared *nod2^-/-^* intestines relative to their wildtype counterparts. However, we also discovered that proliferation rates were equivalent between both genotypes in the absence of commensal microbes (Fig 3F-G), suggesting that NOD2 promotes epithelial proliferation in response to bacterial cues.

To directly test if the canonical NOD2 ligand, MDP, induces intestinal epithelial proliferation via NOD2, we used flow cytometry as an orthogonal, quantitative approach to complement whole-mount imaging and enable unbiased enumeration of cycling intestinal cells following acute ligand exposure.

Using this method, we quantified EdU⁺ cells in dissociated intestines from axenic WT and NOD2-deficient larvae (7 dpf) treated with MDP. We found that MDP treatment more than doubled the number of EdU⁺ cells in axenic WT larvae, establishing it as a potent microbial cue for intestinal growth (Fig 3H). In contrast, the pro-growth effect of MDP was substantially impaired in *nod2^-/-^*larvae with only a minor increase in EdU⁺ cells, demonstrating that the MDP-NOD2 axis links microbial sensing to intestinal stem cell proliferation.

We then assessed proliferation in WT and NOD2-deficient 6-month-old adults using PCNA labeling, which marks cells in S phase of the cell cycle. Here, we once more observed a significant reduction in proliferative PCNA⁺ cells in intestines of NOD2-deficient adults relative to WT controls (Fig 3I, J). Stratification by sex revealed that this reduction was evident in both males and females, with a stronger effect size observed in females (Fig S6A, B). These findings indicate that NOD2-dependent control of epithelial proliferation persists into adulthood and exhibits sex-dependent magnitude. Together, our results establish NOD2 as a regulator of epithelial renewal, linking microbial recognition to epithelial cell proliferation during developmental and adult life stages, and demonstrating an essential role for NOD2 in intestinal growth and homeostasis.

### Loss of NOD2 disrupts epithelial homeostasis by altering cell death and differentiation

While NOD2 is an established microbial sensor, its role in epithelial cell dynamics is less well understood. Our transcriptomic analysis indicated that NOD2 deficiency alters the expression of genes involved in proliferation and cell death across all major secretory epithelial lineages, including goblet cells, enteroendocrine cells, and tuft cells (Fig 2C,4A). As NOD2 dysfunction can impair secretory cell function [44], we determined if NOD2 is required for epithelial cell composition and survival. To characterize the impacts of NOD2 loss on epithelial cell survival and identity in the intestinal epithelial barrier, we quantified apoptotic and Anxa4^+^ cell populations in 6-month-old cohoused WT and *nod2^-/-^*in-crossed siblings (Fig 4B-D). In fish, the 2F11 antibody labels Anxa4^+^ goblet, tuft, and enteroendocrine cells, as well as recently discovered BEST4 cells. We found that *nod2* mutants had a significant increase in TUNEL^+^ cells and a loss of Anxa4^+^ secretory cells relative to WT controls (Fig 4B-D), indicating elevated apoptosis and loss of secretory cell populations in NOD2-deficient guts. Moreover, analysis by sex revealed that both male and female *nod2^-/-^* intestines showed reductions in Anxa4+ secretory cells, an accumulation of Lcp1+ cells, and significantly more TUNEL+ cells compared to WT siblings (Fig S7). These effects were more pronounced in *nod2^-/-^* females, which displayed higher leukocyte infiltration and apoptotic burden relative to *nod2^-/-^*males (Fig S7), suggesting that NOD2 deficiency disrupts epithelial integrity in both sexes but may have a stronger effect on females.

**Fig 4.**
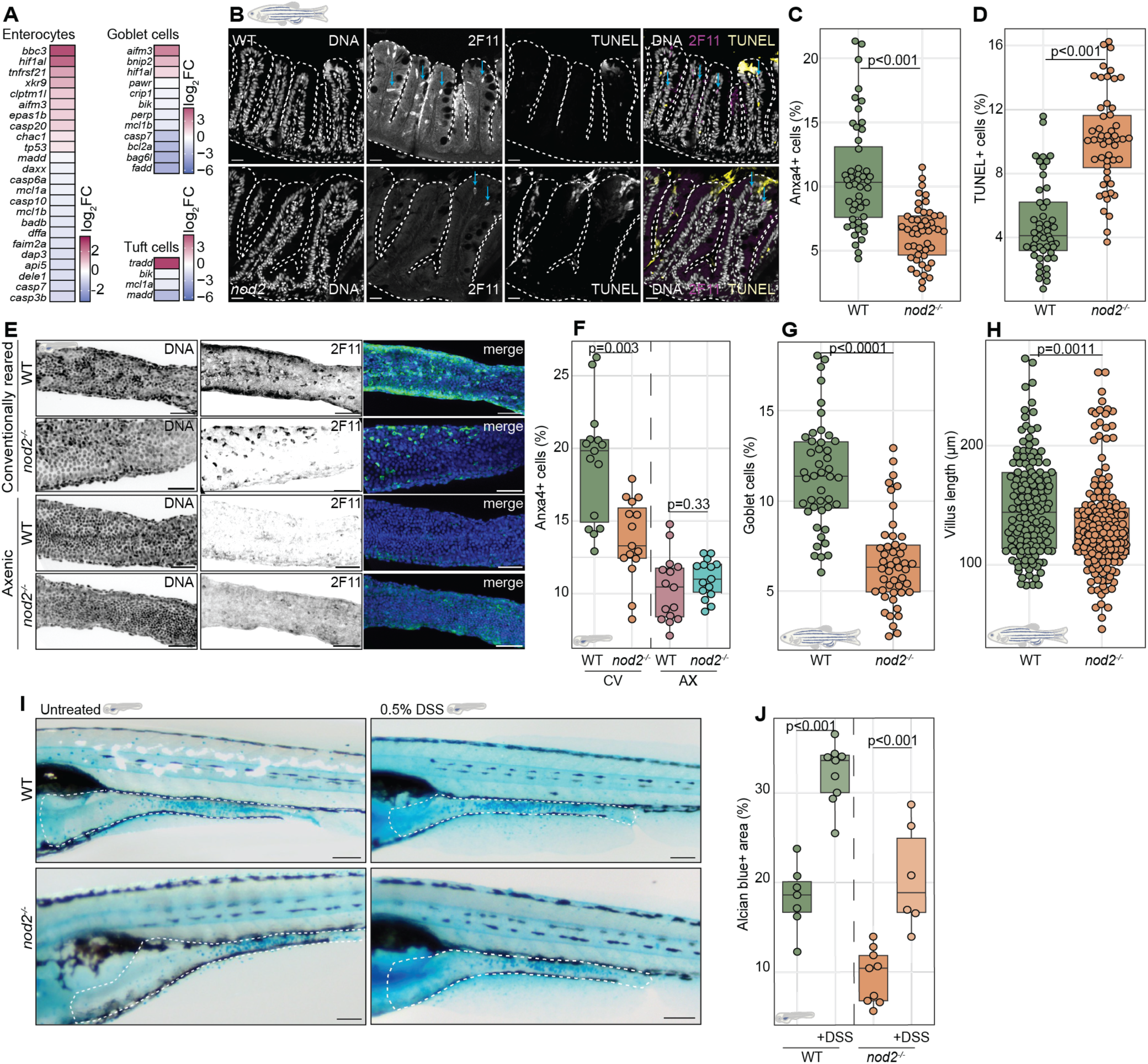
Loss of *nod2* disrupts epithelial homeostasis by altering cell death and cell fate. **A)** Heatmap of cell death gene expression in *nod2^-/-^* ECs, goblet cells, and tuft cells relative to WT controls (for all genes, adjusted p-value ≤ 0.05). **B-D)** Quantification of % Anxa4^+^ cells (C) (blue arrows), and % TUNEL^+^ cells (D) from multiple sections of 6-month-old cohoused adult WT and *nod2^-/-^* intestines. Images are shown in rostral-to-caudal orientation. Each dot represents a single villus measurement from n=5 zebrafish intestines per genotype. **E)** Immunofluorescent images of nuclei (DNA), and Anxa4^+^ (2F11) cells in conventionally reared or axenic WT or *nod2^-/-^* larval intestines (7 dpf), shown in rostral-to-caudal orientation from the mid intestine. In merged false-colored images, DNA is labeled in blue and Anxa4^+^ cells in green. Scale bars = 35μm. **F)** Quantification of the percentage of Anxa4^+^ cells in conventionally-raised (CV) and axenic (AX) larvae. Each dot represents a measurement from a single larval intestine (7 dpf). **G-H)** Quantification of % Alcian-blue+ (goblet cells) (G) and villus length (H) and from multiple sections of 6-month-old cohoused adult WT and *nod2^-/-^* intestines. Each dot represents a single villus measurement from n=5 zebrafish intestines per genotype. **I-J)** Alcian blue staining (I) and quantification of Alcian blue^+^ area (J) in WT, *nod2^-/-^*, WT+DSS, and *nod2^-/-^*+DSS whole mount larvae (7 dpf). Scale bars = 150 μm. P-values were calculated using Mann-Whitney U test or Kruskal-Wallis test. Icons indicate the developmental stage at which the experiment was performed (larval or adult).

**Supplementary Fig 7.**
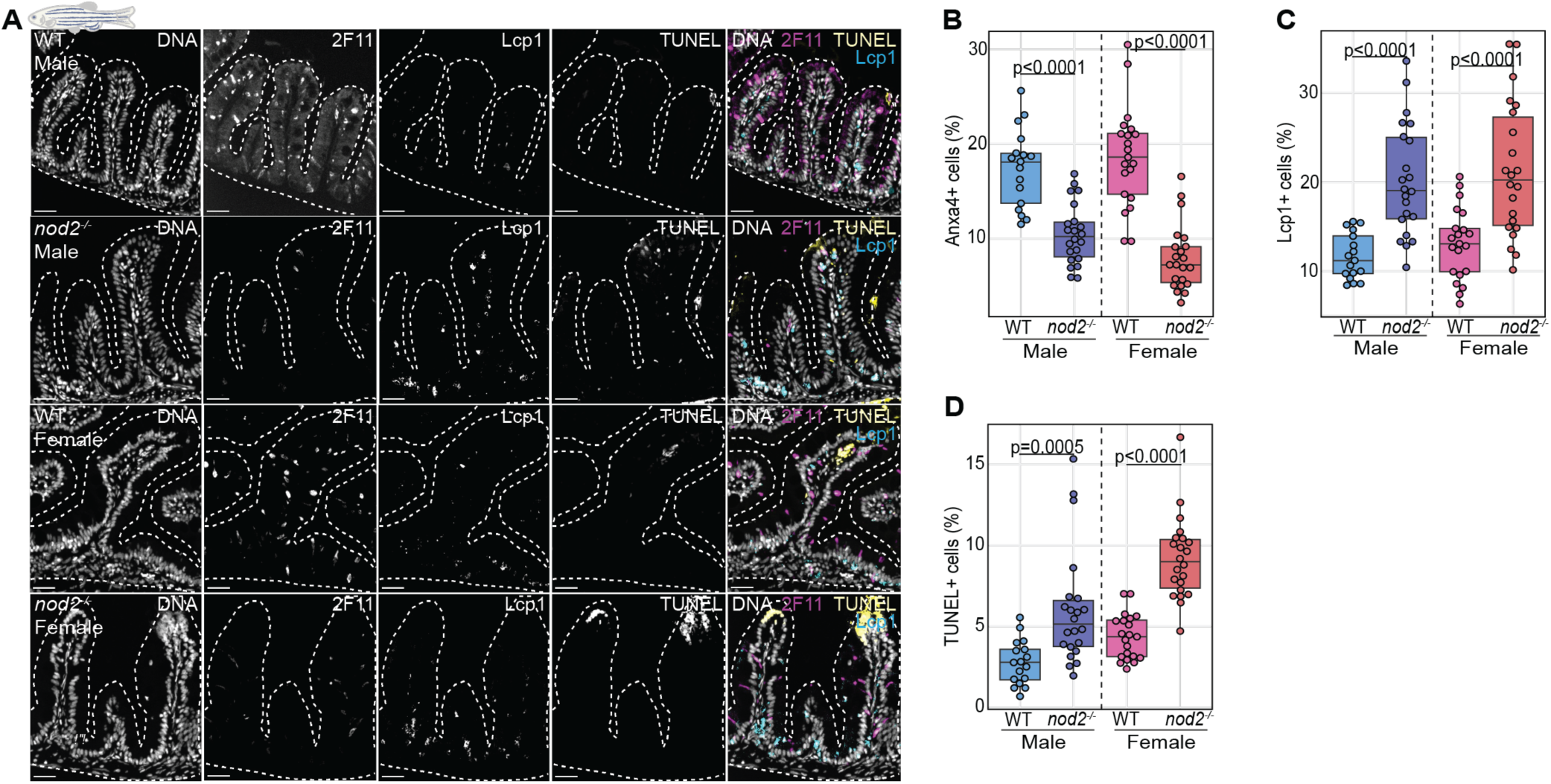
NOD2 deficiency alters epithelial and immune cell populations and increases apoptosis in a sex-dependent manner. **A)** Representative sagittal sections of WT and *nod2^-/-^*intestines (6-month-old, cohoused adults) stained stained for DNA (nuclei), secretory cells (2F11/Anxa4), pan-leukocyte marker (Lcp1), and apoptotic cells (TUNEL). Merged panels show combined staining with outlined villus structures. Images are shown in rostral-to-caudal orientation. **B-D)** Quantification of % Anxa4+ cells (B), % Lcp1+ cells (C), and % TUNEL+ cells (D) from multiple sections of WT and *nod2^-/-^* intestines. Each dot represents a single villus measurement from multiple zebrafish intestines. n = 5 fish per group. Scale bars = 50 μm. P-values were calculated using Mann-Whitney U test or Kruskal-Wallis test. Icon indicates the developmental stage at which the experiment was performed.

**Supplementary Fig 8.**
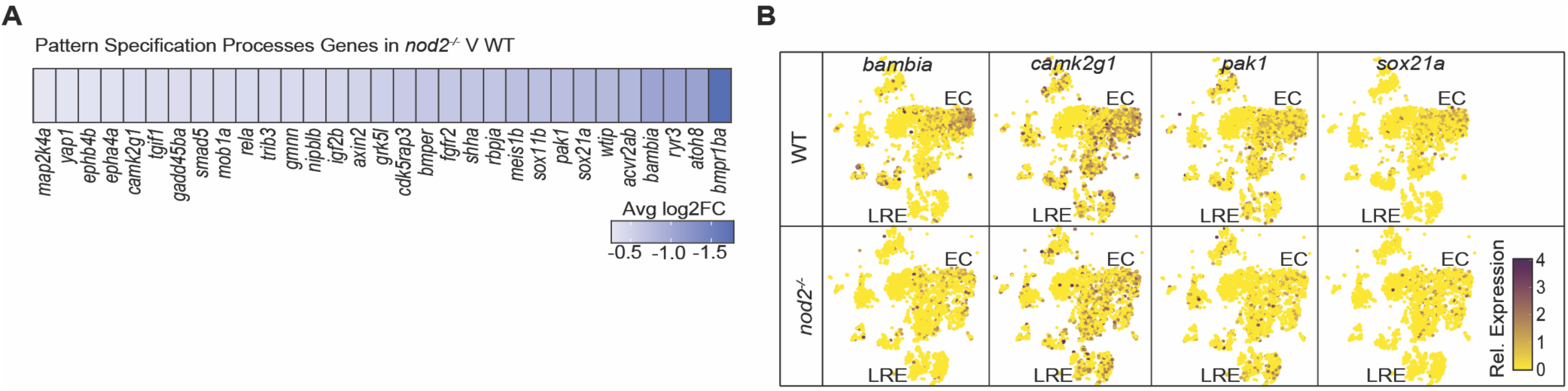
Loss of NOD2 impairs expression of pattern specification genes in the intestinal epithelium. A) Heatmap illustration of representative pattern specification processes gene expression in *nod2^-/-^* versus WT adult intestinal epithelium (for all genes, adjusted p-value ≤ 0.05). B) Feature plots showing relative expression of select patterning genes (*bambia*, *camk2g1*, *pak1*, *sox21a*) in WT and *nod2^-/-^* intestinal epithelium. Expression is reduced across enterocyte (EC) and lower right epithelial (LRE) clusters in *nod2^-/-^* intestines, indicating broad loss of epithelial patterning cues.

Beyond cell survival, epithelial architecture depends on transcriptional patterning programs that establish the cellular heterogeneity required for intestinal function. Our differential gene expression analysis uncovered downregulated expression of key patterning genes in *nod2^-/-^* intestines, including *map2k4a*, *epb41l4b*, *camk2g1*, *pak1*, and *sox21a* (Fig S8A, B), suggesting that NOD2 modifies genetic programs that regulate spatial identity and cell fate. To test roles for NOD2 in epithelial patterning, we quantified Anxa4^+^ secretory cells in intestines of WT and *nod2^-/-^* larvae (7 dpf) raised under either conventional or axenic conditions. Under conventional conditions, *nod2^-/-^* larvae exhibited a significant reduction in Anxa4^+^ cells compared to WT siblings (Fig 4E, F). This reduction was not observed in axenic *nod2^-/-^*larvae, suggesting that commensal microbes typically engage NOD2 to promote maturation of secretory lineages in the gut. Supporting this, we discovered that adult *nod2^-/-^* intestines also exhibited shorter villi and fewer goblet cells (Fig 4G, H), reflecting impaired secretory lineage differentiation, and mirroring patient data that associate NOD2 mutations with goblet cell loss [45, 46].

To directly assess the impact of NOD2 deficiency on goblet cell-associated mucus output, we performed whole-mount Alcian blue staining of intestinal mucus in wildtype and *nod2* mutant larvae (7 dpf). We performed the quantification using a fixed, pre-defined color threshold applied uniformly across all images with blinded analysis to prevent observer bias. Because Alcian blue labels mucins rather than goblet cells themselves, this assay reports mucin-positive area and not goblet cell number or density. We found that *nod2* mutants displayed a significant reduction in Alcian blue^+^ area under homeostatic conditions (Fig 4I, J), indicating reduced goblet cell function. Treatment with dextran sulfate sodium (DSS), a colitogenic polysaccharide that induces epithelial damage and inflammation in vertebrates, led to an expansion of the Alcian blue^+^ mucin signal in both genotypes, with stained area approximately doubling in each. While the relative increase was similar, mucin-positive area remained lower in *nod2^-/-^*larvae, suggesting that epithelial stress can partially rescue goblet cell function through NOD2-independent mechanisms, though the response remains blunted in the absence of NOD2.

Combined, our observations point to a critical role for NOD2 in maintaining epithelial homeostasis, but raise a key mechanistic question: what is the underlying mechanism that contributes to the epithelial phenotypes observed in *nod2^-/-^*mutants?

### NOD2 deficiency enhances estrogen signaling in the intestinal epithelium

To investigate the molecular pathways responsible for epithelial disruption in *nod2^-/-^* intestines, we examined our transcriptomic data for signatures of impaired signaling in NOD2-deficient adult intestines and identified enrichment of estrogen response pathways as the most prominently affected (Fig 5A-B). The zebrafish genome encodes *esr1*, and *esr2*, orthologs of mammalian ERα and ERβ, respectively. In fish, a genome duplication event led to evolution of *esr2a* and *esr2b* paralogs from the ancestral *esr2* gene. We observed enriched expression of *esr2b* specifically in IECs (Fig 5C), and widespread upregulation of estrogen-response genes, including *esr2b* and multiple vitellogenins (*vtg1-vtg7*), across epithelial lineages in *nod2^-/-^* intestines compared to their wildtype sibling controls (Fig 5D, E). Notably, estrogen-response genes were also elevated in immune and stromal subsets of *nod2^-/-^*intestines (Fig 5F, G), suggesting that NOD2 deficiency broadly impacts estrogenic signaling across the intestinal cellular landscape. To validate the estrogen-related transcriptional changes observed in our scRNA-seq data, we performed qPCR for *vtg1*, *vtg2*, and *vtg4* in 6-month-old co-housed WT and *nod2^-/-^* zebrafish intestines, separated by sex (Fig 5H, I). We found that, in males, *vtg1*, *vtg2* and *vtg4* levels remained unchanged between WT and *nod2^-/-^* (Fig 5H). In contrast, *nod2^-/-^* females exhibited significant increases in *vtg* gene expression compared to WT controls (Fig 5I), indicating that NOD2 deficiency enhances estrogen-responsive gene expression in a sex-specific manner.

**Fig 5.**
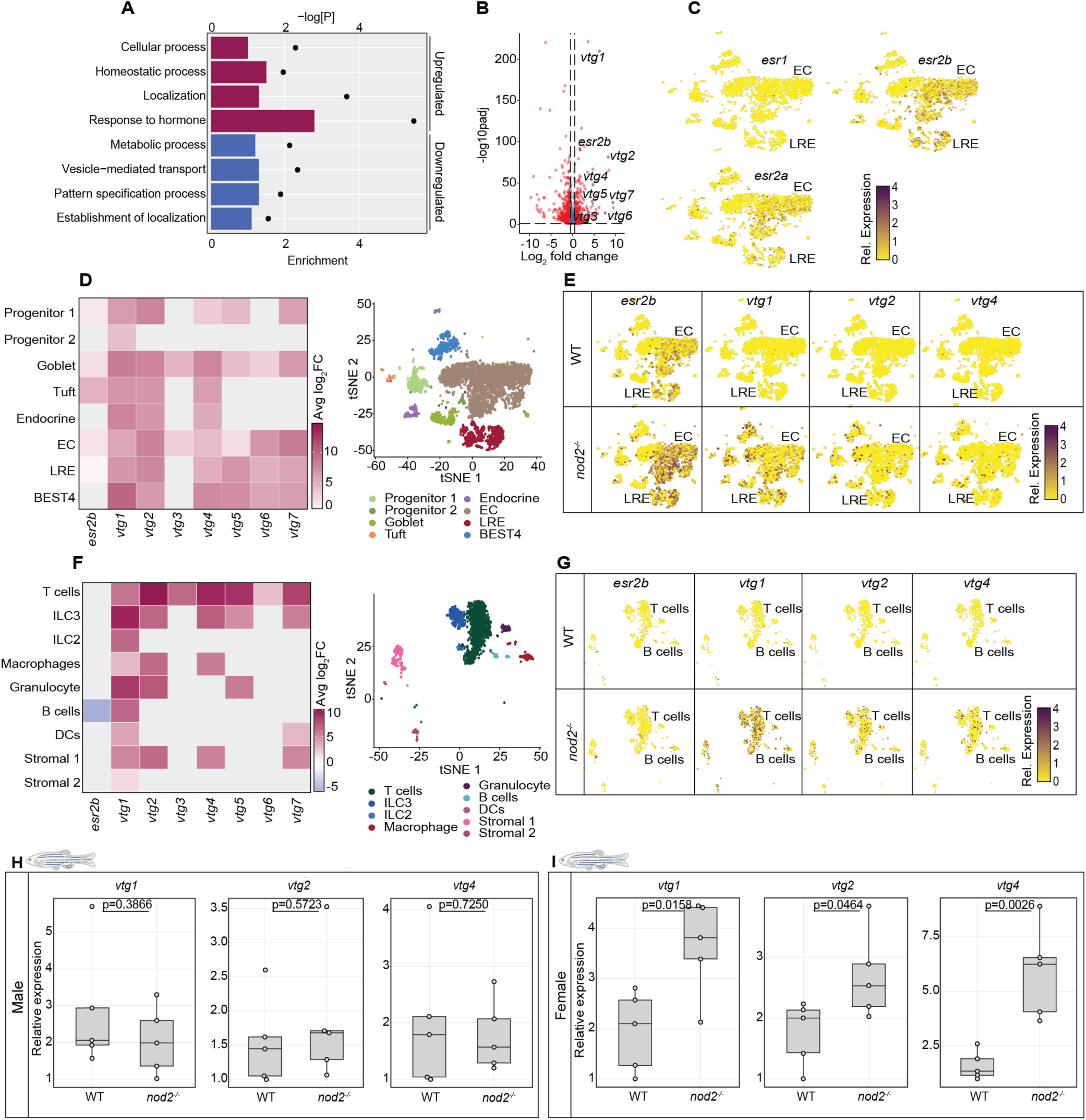
NOD2-deficiency significantly increased the intestinal estrogen response. **A)** Top four up, or downregulated gene ontology terms in *nod2^-/-^* epithelium relative to WT controls. Relative enrichment score for each term is indicated by bar length and negative log P-values are shown with closed circles. **B)** Volcano plot of integrated dataset DE genes in *nod2^-/-^* epithelium relative to WT controls. Significance values represented on the y-axis and relative fold change of gene expression on the x axis. Only genes with log2 expression changes greater than 0.25 are shown. Genes listed as response to hormone are indicated. **C)** Feature plots showing relative expression of the estrogen receptors *esr1*, *esr2a*, and *esr2b* in intestines of WT fish. Enterocytes (EC) and Lysosome-Rich Enterocytes (LRE) are labeled for orientation. **D)** Heatmap (left) depicting relative changes in estrogen-response gene expression, including *esr2b*, in *nod2^-/-^* versus WT epithelium (for all genes, adjusted *p* ≤ 0.05), and tSNE plot (right) of epithelial cell type annotations. **E)** Feature plots showing expression of *esr2b*, *vtg1*, *vtg2*, and *vtg4* in WT and *nod2^-/-^* datasets across epithelial lineages. **F)** Heatmap (left) and tSNE plot (right) of immune subsets showing relative changes in estrogen-response gene expression in *nod2^-/-^*versus WT. **G)** Feature plots showing *esr2b*, *vtg1*, *vtg2*, and *vtg4* expression in WT and *nod2^-/-^* immune subsets. **H-I)** qPCR analysis of the relative expression of estrogen-sensitive vitellogenin genes (*vtg1*, *vtg2*, and *vtg4*) in WT and *nod2^-/-^* adult zebrafish intestines, separated by sex (6-month-old, cohoused adults). P-values were calculated using Mann-Whitney U test or Kruskal-Wallis test. Icon indicates the developmental stage at which the experiment was performed.

### Estrogen signaling modulates intestinal epithelial proliferation, goblet cell abundance, and leukocyte recruitment through a NOD2-dependent mechanism

Given the links between estrogen signaling and epithelial cell proliferation and survival [47–49], we asked if estrogen and NOD2 interact to control intestinal cell fates. To this end, we analyzed epithelial function in WT and NOD2-deficient larvae (7 dpf) that we treated with 17β-estradiol (E2), or the selective estrogen receptor modulator, tamoxifen, which acts as a potent estrogen receptor antagonist in larval zebrafish. As larvae are estrogen-sensitive hermaphrodites [50], they provide a unique opportunity to examine sex-independent effects of estrogen on gut physiology. To test if interactions between estrogen and NOD2 modify intestinal renewal, we first quantified intestinal epithelial proliferation in wildtype and *nod2^-/-^* larvae that we treated with E2, or the vehicle control DMSO, for 24 hours. We discovered that E2 treatment reduced the number of proliferative cells by over 50% in wildtype larvae but had negligible impacts on proliferation in NOD2-deficient counterparts (Fig 6A, B). Notably, pre-treatment with tamoxifen [51] had the opposing effect. Tamoxifen exposure significantly elevated intestinal proliferation rates in NOD2-deficient larvae without discernible impacts on wildtype controls (Fig 6C, D).

**Fig. 6.**
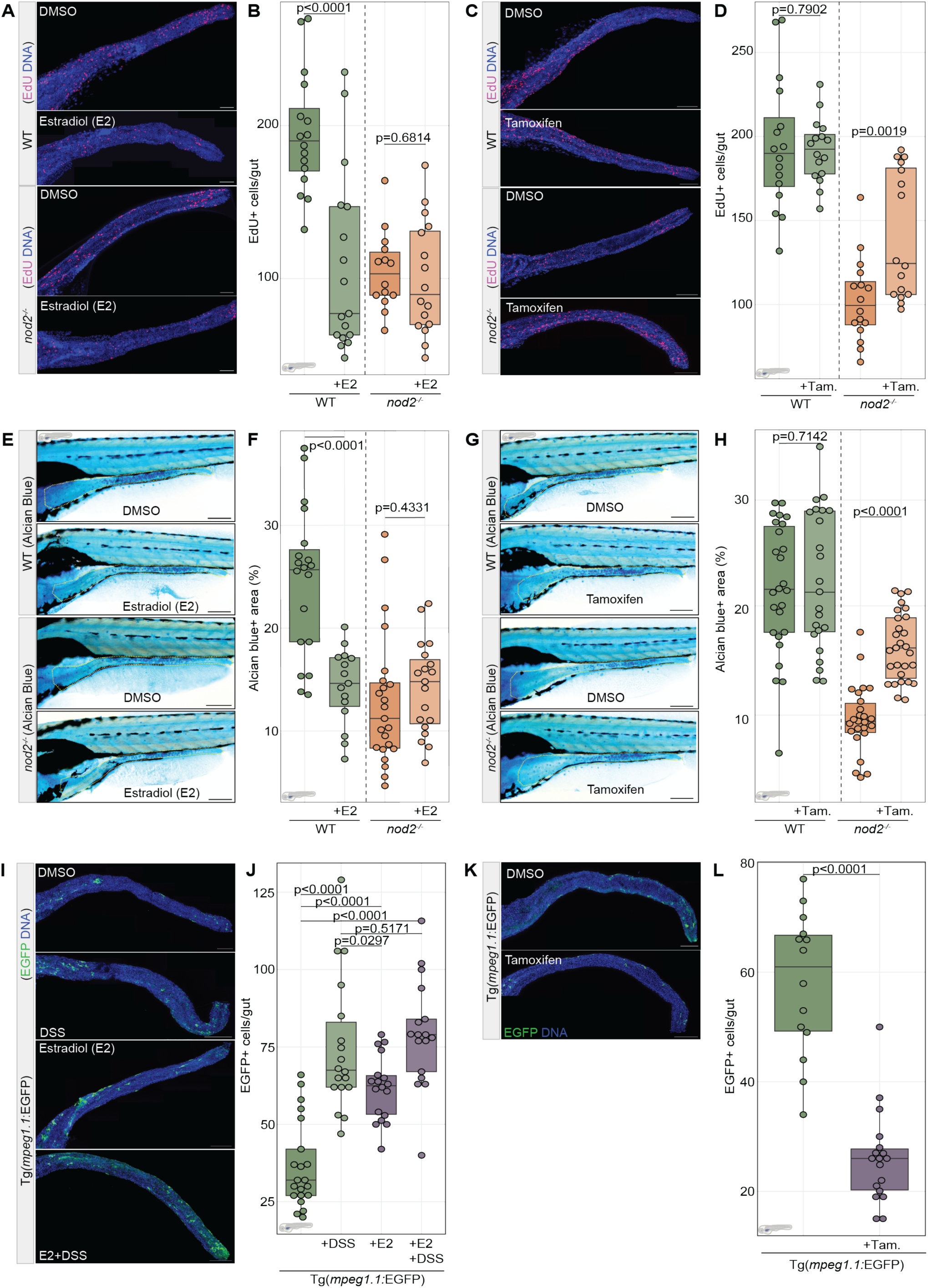
NOD2-dependent estrogen signaling modulates intestinal epithelial proliferation, goblet cell abundance, and macrophage recruitment. **A)** Immunofluorescence of whole-mount WT and *nod2^-/-^* larval intestines (7 dpf) treated with DMSO (vehicle control) or 25 ng/μL estradiol (E2). In merged false-colored images, DNA is labeled in blue and EdU^+^ proliferating cells in magenta. Scale bars = 150 μm. **B)** Quantification of EdU^+^ cells per gut. Each dot represents one fish. Estrogen significantly reduced proliferation in WT larvae but not in *nod2^-/-^.* C) Immunofluorescence of WT and *nod2^-/-^* larval intestines (7 dpf) treated with DMSO or 1.5 ng/μL Tamoxifen (Tam). DNA is labeled in blue and EdU+ cells in magenta. Scale bars = 150 μm. **D)** Quantification of EdU^+^ cells per gut. Tamoxifen increased proliferation in *nod2^-/-^*larvae but had no effect in WT. **E-F)** Alcian blue staining (E) and quantification of Alcian blue^+^ area (F) in WT and *nod2^-/-^* larvae (7 dpf) treated with DMSO or E2. E2 significantly reduced goblet cell area in WT, but not in *nod2^-/-^* larvae. **G-H)** Alcian blue staining (G) and quantification (H) of goblet cell area in WT and *nod2^-/-^* larvae (7 dpf) treated with DMSO or tamoxifen. Tamoxifen significantly increased goblet cell abundance in *nod2^-/-^* larvae. **I)** Immunofluorescence of whole-mount *Tg(mpeg1*:EGFP) larval intestines (7 dpf; shown in rostral-to-caudal orientation from the base of the intestinal bulb to the cloaca) pre-treated with DMSO or 25 ng/μL E2 for 24hrs and the exposed to +/-0.5% DSS for 24hrs. In merged false-colored images, DNA is labeled in blue and EGFP+ macrophages in green. Scale bars = 150 μm. **J)** Quantification of EGFP+ cells per gut from *Tg(mpeg1*:EGFP) larvae. Each dot represents an individual intestine. **K-L)** Representative images (K) and quantification (L) of EGFP+ macrophages in *Tg(mpeg1*:EGFP) larvae (7 dpf; shown in rostral-to-caudal orientation from the base of the intestinal bulb to the cloaca) treated with DMSO or Tam. Tam. significantly reduced macrophage numbers in *Tg(mpeg1*:EGFP) intestines. Each dot represents an individual intestine. P-values were calculated using the Mann-Whitney U test. Icon indicates the developmental stage at which the experiment was performed.

Looking at goblet cell function, we detected similar interactions between NOD2 and estrogen. Exposure to E2 significantly reduced the Alcian blue^+^ area of wildtype larval intestines (7 dpf) but had no effect on Alcian blue levels in *nod2^-/-^* intestines. (Fig 6E, F), suggesting that epithelial responses to estrogen are maximally engaged in the absence of NOD2. Finally, we found that tamoxifen treatment nearly doubled the intestinal Alcian blue-positive area of *nod2^-/-^* intestines, with no discernible effects on WT siblings (Fig 6G, H).

To determine if estrogen-NOD2 interactions also influence the intestinal accumulation of gut-associated phagocytes, we analyzed macrophage populations in intestines of *Tg(mpeg1.1*:EGFP) larvae that we treated with E2 or tamoxifen for 24hrs or 6hrs, respectively (7 dpf). Supporting effects of E2 on mucosal defenses, we discovered that E2 treatment more than doubled the number of gut-associated macrophages in wildtype larvae, matching levels found after DSS exposure, whereas tamoxifen exposure significantly reduced macrophage accumulation (Fig 6I-L). In sum, our work shows that treatment of wildtype larvae with E2 replicates the proliferative and barrier defects noted in NOD2-deficient larvae, while tamoxifen exposure partially rescues barrier defects in *nod2^-/-^* intestines, indicating a role for the NOD2-estrogen axis in the maintenance of intestinal epithelial homeostasis.

### Estrogen signaling impairs epithelial recovery in a NOD2-dependent manner

Finally, to assess the consequences of estrogen-NOD2 interactions during colitogenic challenges, we monitored survival rates of wildtype and NOD2-deficient larvae that we exposed to DSS. Kaplan-Meier survival analysis revealed a significant reduction in survival for *nod2^-/-^*larvae following DSS exposure compared to WT controls (Fig 7A), consistent with previous work [31] showing increased sensitivity of NOD2-deficient zebrafish to epithelial injury.

**Fig 7.**
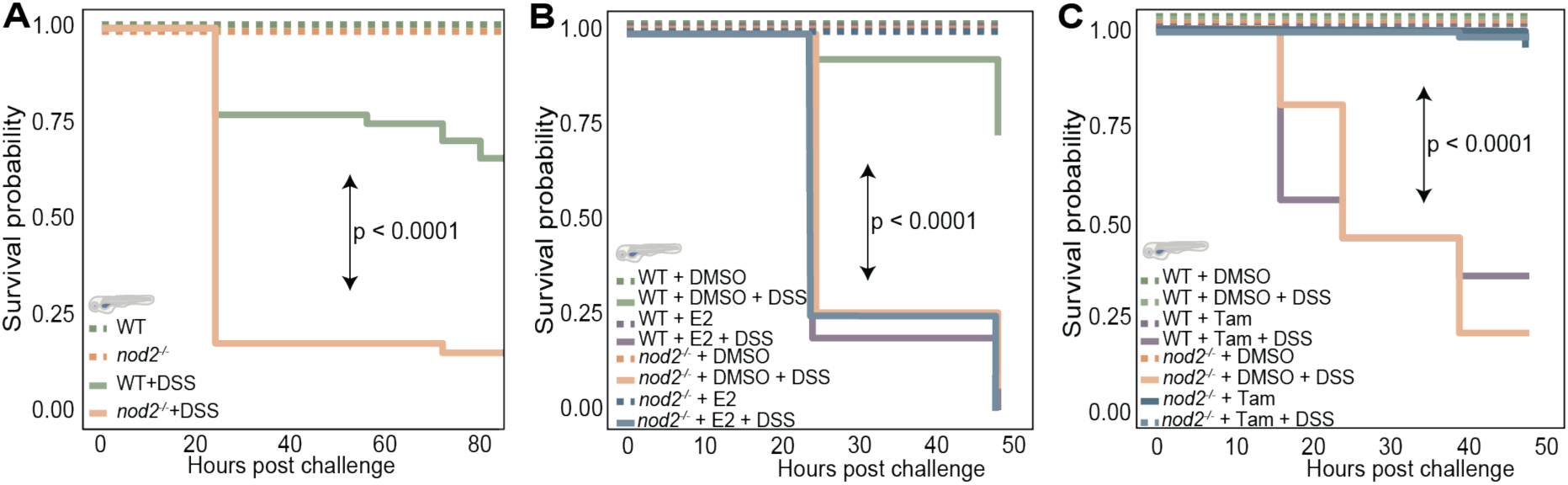
Estrogen modulates the survivability of DSS-treated larvae in a NOD2-dependent manner. **A)** Kaplan–Meier survival curves comparing WT and *nod2^-/-^* larvae exposed to 0.5% DSS beginning at 4 dpf, with n = 20 larvae per group. *nod2^-/-^* larvae exhibit significantly reduced survival following DSS-induced intestinal injury relative to WT siblings, indicating heightened sensitivity to epithelial damage in the absence of NOD2. **B)** Survival analysis of WT and *nod2^-/-^*larvae pretreated with either DMSO or 17β-estradiol (E2) beginning at 4 dpf, followed by 0.5% DSS exposure at 5 dpf. P-values were determined using the log-rank (Mantel-Cox) test. **C)** Survival analysis of WT and *nod2^-/-^* larvae with 6hr pretreatment with DMSO or tamoxifen (Tam.) beginning at 5 dpf, followed by 0.5% DSS exposure at 6 dpf. Tamoxifen significantly improved survival in *nod2^-/-^* larvae compared to vehicle-treated controls. Icon indicates the developmental stage at which the experiment was performed.

We next asked if estrogen modulates intestinal resilience in a NOD2-dependent manner. To this end, we treated WT and *nod2^-/-^*larvae with E2 prior for 24hrs to a DSS challenge. As anticipated, we found that *nod2^-/-^* larvae exhibited poor survival regardless of E2 treatment status, and we found that wildtype larvae effectively survived an acute DSS challenge, confirming a role for NOD2 in regulating host responses to a pro-inflammatory challenge (Fig 7B). Strikingly, we discovered that E2-treated WT larvae phenocopied the heightened mortality observed in *nod2^-/-^* animals, indicating that estrogen impairs epithelial recovery in a NOD2-dependent mechanism (Fig 7B).

Finally, we tested if tamoxifen pretreatment restores resilience to DSS exposure in *nod2* mutant larvae. Remarkably, tamoxifen significantly improved survival in *nod2^-/-^* larvae (Fig 7C), indicating that attenuation of estrogen signaling rescues NOD2-associated defects in epithelial recovery. However, we also found, unexpectedly, that WT larvae displayed enhanced sensitivity to DSS following pre-treatment with tamoxifen. We speculate that this opposing response suggests that the effects of estrogen receptor blockade on intestinal integrity depend on the underlying inflammatory context and basal NOD2 activity, rather than acting as a uniform protective mechanism.

In sum, our work shows that loss of NOD2 significantly elevates E2 signaling in the intestine, that E2 exposure is sufficient to recapitulate several key NOD2-deficiency phenotypes in zebrafish larvae, and that pharmacological attenuation of E2 signaling restores epithelial homeostasis in NOD2-deficient fish, indicating that estrogen and NOD2 act in opposing directions to regulate intestinal homeostasis.

## DISCUSSION

Loss-of-function *NOD2* alleles are the single greatest genetic risk factor for Crohn’s disease. However, we only have a partial appreciation of the consequences of NOD2 deficiency for intestinal epithelial cell dynamics prior to the manifestation of symptoms. We consider this an important gap in our understanding of the factors that promote a chronic inflammatory illness that continues to grow in frequency.

In this study, we used zebrafish to determine how NOD2 deficiency impacts homeostatic maintenance of the intestinal barrier. Fish guts are remarkably like their mammalian counterparts in terms of developmental regulation and cell type composition [23–29], and NOD2-deficient fish replicate key features of Crohn’s disease [30, 31]. Early zebrafish work established conserved Nod pathway expression and antibacterial function in larvae using splice-blocking morpholinos targeting the *nod2* exon 1 donor site [30]. In that study, *nod2* knockdown did not grossly disrupt larval development but reduced host control of systemic *Salmonella* infection and was associated with decreased intestinal epithelial expression of *duox*, consistent with a conserved role for Nod signaling in mucosal antibacterial defense. Work with a CRISPR/Cas9-generated *nod2* mutant reported Crohn’s disease-relevant intestinal phenotypes, including altered gut morphology, leukocyte accumulation, and heightened sensitivity to DSS-induced epithelial injury, with additional outcomes such as fibrosis and epithelial hypertrophy reported under inflammatory challenge [31]. Our line was generated independently using a CRISPR-induced frameshift allele upstream of the leucine-rich repeat domain and is likewise predicted to severely impair NOD2 function. Together, the convergence of *nod2*-dependent phenotypes across transient knockdown and independent stable mutant alleles supports a conserved requirement for NOD2 in intestinal immunity and barrier homeostasis. Consistent with these studies, we found that, like mouse models, NOD2 deficiency alone does not lead to overt inflammatory disease in fish, and that *nod2* genetic status has minimal impacts on microbiota composition, particularly in the context of co-housing [15], emphasizing the utility of fish as a genetically accessible model of NOD2 deficiency.

In humans, Crohn’s disease is not typically associated with complete NOD2 deletion but instead with common coding variants, including frameshift and missense alleles, that reduce or alter NOD2 signaling capacity [2,3,5]. Several of the best-characterized risk alleles are functionally loss-of-function with respect to MDP sensing and downstream NF-κB activation, although they may retain partial or context-dependent activity [8,12,13]. Rodent models similarly include both null and hypomorphic Nod2 alleles that produce overlapping barrier and inflammatory phenotypes [17–19,33]. Our zebrafish null model therefore represents the severe end of the NOD2 functional spectrum and is well suited to define core NOD2-dependent homeostatic mechanisms, while human disease likely reflects partial impairment layered onto additional genetic and environmental modifiers.

Consistent with established roles for NOD2 in immune regulation, we observed immune dysregulation in hematopoietic, epithelial, and stromal compartments of *nod2* mutants, including altered expression of inflammatory cytokines and autophagy pathway components that are widely linked to CD. However, we also found that NOD2 deficiency affected epithelial renewal and organization in an estrogen-sensitive manner. These findings add to our mechanistic understanding of NOD2 gut function and raise unanswered questions for future studies.

How does NOD2 regulate intestinal epithelial cell dynamics? Working with co-housed, wildtype and *nod2* mutant siblings derived from a heterozygous in-cross, we demonstrated that NOD2 deficiency leads to intestinal shortening, increased sensitivity to DSS-induced damage, and reduced cell survival. Our results match findings from *NOD2* mutant mice and patient data, which show altered intestinal permeability [8], and increased susceptibility to inflammatory injury [18,19]. However, we do not know how loss of NOD2 translates to barrier defects. Our transcriptional data revealed widespread dysregulation of genes linked to growth and differentiation, and our quantitative work showed that NOD2 is necessary for epithelial proliferation and maturation. However, we also discovered shifts in the transcriptional landscape of genes required for cell survival, and we found that NOD2 deficiency enhanced the incidence of epithelial cell death, especially within *nod2^-/-^* females. Therefore, future studies using lineage tracing and cell-fate labeling strategies are required to resolve the extent to which cell proliferation, differentiation and survival accounts for barrier defects in *nod2* mutants.

How do NOD2 and estrogen interact to organize the epithelial barrier? One of the most striking findings from our transcriptomic analysis was a pronounced increase of estrogen signaling in NOD2-deficient guts. Confirming links between NOD2 deficiency, estrogen signaling, and gut function, we showed that exposure to E2 enhances intestinal accumulation of macrophages, attenuates epithelial proliferation, and diminishes generation of goblet cells in wildtype larvae, whereas tamoxifen reduces macrophage accumulation, restores epithelial proliferation, and increases goblet cell numbers in NOD2-deficient larvae. Survival assays extended these observations by showing that E2-treated wildtype larvae phenocopy the DSS sensitivity of *nod2* mutants, while tamoxifen improves survival in DSS-treated *nod2^-/-^* mutants. Combined, these findings indicate that estrogen is a physiologically relevant modifier of NOD2 function in the gut. As a caveat, we found that tamoxifen treatment had the unexpected effect of rendering wildtype larvae more sensitive to DSS, indicating a need for future examination of interactions between NOD2 and estrogen in pro-inflammatory contexts.

Further supporting a mechanistic link between NOD2 and estrogen signaling bazedoxifene reduces fibrosis, prevents epithelial hypertrophy, and suppresses pro-inflammatory cytokine expression in *nod2* mutants treated with DSS [31]. Although that work employed bazedoxifene as gp130 antagonist, we note that bazedoxifene is primarily known as a selective estrogen response modulator, widely prescribed for managing postmenopausal osteoporosis in women. Consistent with a broader role for estrogen receptor modulation in host-microbe interactions, a recent zebrafish study demonstrated that tamoxifen treatment alters susceptibility and immune responses to *Mycobacterium marinum* infection [53], further supporting that estrogen pathway modulators can reshape mucosal and inflammatory outcomes *in vivo*. Together with growing literature linking estrogen signaling to epithelial barrier regulation and intestinal inflammation, these findings place our NOD2-estrogen interaction within an emerging cross-species framework connecting hormone signaling to gut immune homeostasis.

Commonly associated with development of female secondary sexual characteristics, estrogen has non-gonadal roles in male and female somatic tissue, including in IBD [54]. Epidemiological data showed that women have a greater incidence of Crohn’s disease and experience more extraintestinal comorbidities than men [55–57]. Confirming a causal relationship between estrogen and Crohn’s disease, oral contraceptive pill usage significantly increases disease risk for women [58]. In the gut, estrogen engages the nuclear estrogen receptors ERα and ERβ to control expression of target genes and interacts with the membrane-bound G protein-coupled estrogen receptor, GPER, to modify cellular functions at the post-transcriptional level [59]. In our intestinal single-cell RNA-seq dataset, *gper1* expression was not detectably enriched above background in epithelial or immune cell populations in both WT and *nod2* mutants, suggesting that nuclear estrogen receptor signaling is the dominant estrogen-responsive pathway captured in our gut analyses, yet a contribution from membrane estrogen receptors cannot be excluded in other contexts. Mechanistically, animal and tissue culture work indicate that estrogen primarily signals through ERβ to arrest proliferation and enhance cell death [47–49], matching phenotypes noted in the current study, and leading us to hypothesize that estrogen promotes an accumulation of barrier defects in *nod2* mutants. However, the outcome of estrogen signaling in the intestine also depends on the relative abundance and activity of ERa and ERb. Shifts in the ERα : ERβ ratio can reverse epithelial responses and influence inflammatory susceptibility [60,61]. Recent mammalian work demonstrated that epithelial-restricted Erα, rather than Erβ, is the primary determinant of female-biased protection from colitis, establishing epithelial estrogen receptor signaling as a key driver of sex differences in mucosal resilience [62,63]. In contrast, our zebrafish data revealed upregulation of *esr2b*, the ERβ-related paralog, in *nod2^-/-^* intestinal epithelial cells, suggesting that NOD2 deficiency could trigger a receptor-specific estrogenic compensation program that differs from the ERα-dominant response described in mammals.

While our data support a functional interaction between NOD2 and estrogen, the molecular link remains unresolved. It is plausible that NOD2 influences estrogen receptor stability, localization, or downstream transcriptional activity, either directly or through intermediates such as MAPK or PI3K signaling, both of which are modulated by NOD2 and have known interactions with estrogen pathways [64–66]. Alternatively, loss of NOD2 may lead to compensatory estrogen receptor activation in response to epithelial stress. Investigating these mechanistic connections will be an important area for future work. What about sex? Our study design reflects the chronological trajectory of discovery. Initial transcriptomic and phenotypic analyses were conducted using mixed-sex cohorts to define core NOD2-dependent intestinal phenotypes. These early experiments were designed to identify genotype-driven effects shared across animals rather than sex-specific responses. However, combined with prior work demonstrating sex-specific interactions between estrogen and inflammatory disease in the gut [62,63], our transcriptomic data revealed a strong estrogen-responsive signature in *nod2* mutants, indicating that sex and hormonal context can modify downstream outcomes. This prompted targeted follow-up analyses with sex recorded and evaluated explicitly. Sex-stratified measurements of epithelial proliferation, apoptosis, secretory lineages, and immune cell accumulation demonstrate that NOD2 regulates intestinal homeostasis in both sexes, while the magnitude of several phenotypes is greater in females. We therefore interpret NOD2 as a regulator of shared epithelial and immune programs whose phenotypic expression is modulated by hormonal environment rather than restricted to a single sex. However, defining the full scope of sex-specific transcriptional and functional responses, including receptor-specific estrogen pathway effects, will require dedicated, sex-balanced experimental designs and represents an important direction for ongoing work beyond the scope of the present manuscript.

We feel our findings have particular relevance to CD. The discovery that NOD2 interacts with estrogen offers a mechanistic insight into how its loss might impair epithelial barrier function. Estrogen has been implicated in the sex-dependent pathophysiology of CD with notable age-dependent differences in onset and severity. In pediatric cases, girls often present earlier and more severely than boys [67]. In adults, women have lower CD incidence during reproductive years, but risk rises post-menopause [68–74]. Additional clinical data shows that symptoms fluctuate with menstrual cycles [75–77], and that irregular menstruation often precedes diagnosis [78]. Our finding that *nod2^-/-^* female fish exhibit exaggerated estrogen responses may help explain how the hormonal environment interacts with genetic susceptibility to shape disease risk.

Future studies should investigate whether this regulatory axis operates in other mucosal tissues or during hormonal transitions such as puberty, pregnancy, or menopause. Additionally, given the increasing interest in estrogen modulators as therapeutic agents, our results suggest that NOD2 genotype may influence responses to hormone-based therapies and should be considered in precision medicine approaches to IBD.

## LIMITATIONS

While our data support a NOD2-estrogen axis, several constraints should be noted. Tamoxifen combined with DSS caused severe lethality in WT larvae, preventing leukocyte quantification in *Tg(mpeg1.1*:EGFP) fish under these conditions. This likely reflects a threshold effect, where concurrent ER antagonism and epithelial injury exceed the tolerance of the WT gut, resulting in barrier collapse rather than a graded response. Consequently, immune recruitment under tamoxifen + DSS could not be assessed and will require dose titration or staggered treatment strategies.

As with all larval pharmacology, E2 and tamoxifen may engage additional target pathways, and the responsible ER isoform remains unconfirmed. Our transcriptional data support involvement of nuclear estrogen receptors, particularly *esr2b*, but do not exclude contributions from membrane-associated estrogen signaling pathways. In our intestinal single-cell dataset, *gper1* transcripts were not detected in any gut cell population in either WT or *nod2* mutants, preventing assessment of its contribution in this context. Resolving receptor-specific mechanisms will require targeted genetic or pharmacologic interrogation in future studies.

In addition, although the *nod2* mutant phenotypes are robust and reproducible across assays, we cannot formally exclude the possibility that the allele functions as a hypomorph rather than a complete null. The small residual responses observed in *nod2* mutants following MDP stimulation are consistent with this possibility. However, such compensation would be expected to attenuate rather than generate the phenotypes we report, and therefore would not alter the overall interpretation of a NOD2-dependent role in epithelial and immune regulation.

Lastly, while several key adult phenotypes are now presented in sex-stratified form, not all experiments were originally powered for sex-specific analysis, as early discovery-stage datasets were designed to identify shared genotype-dependent effects. Future work will incorporate prospectively balanced and fully sex-stratified designs to resolve hormone-genotype interactions at higher resolution and to distinguish baseline NOD2-dependent programs from sex-hormone-dependent modifiers.

## MATERIALS AND METHODS

### Zebrafish strains and maintenance

All zebrafish experiments were performed at the University of Alberta using protocols approved by the University’s Animal Care & Use Committees, Biosciences and Health Sciences (#3032), operating under the guidelines of the Canadian Council of Animal Care. Wild-type (TL), *nod2^-/-^* knockout zebrafish and *Tg(mep1.1*:EGFP) (four to nine months-old; approx. equal sex distribution) were reared at 29°C in tank water (Instant Ocean Sea Salt dissolved in reverse osmosis water maintained at a conductivity of 1000μS and pH buffered to pH 7.0 with sodium bicarbonate) under a 14h/10h light/dark cycle using standard zebrafish husbandry protocols. Feeding regimens were age specific. From 6-14 days post-fertilization (dpf), larvae were fed GEMMA Micro 75 twice daily and live rotifers once daily. From 15-28 dpf, larvae received a 1:1 mixture of GEMMA Micro 75 and GEMMA Micro 150 twice daily, along with live rotifers once daily. Juvenile fish (29 dpf to 13 weeks) were fed GEMMA Micro 300 twice daily and live rotifers once daily. Adult zebrafish were maintained on a once-daily diet of GEMMA Micro 300 and live rotifers. For larval analysis, breeding tanks were set up overnight with 1 male and 1 female separated by a divider until morning. Fish were bred for 1 hour, then embryos were collected, rinsed gently with facility water, and transferred to a petri dish (fisherbrand) with 30 mL embryo media (EM) and 30 embryos per dish. Embryos were raised at 29°C under a 14 hour/ 10 hour light/ dark cycle until 7 dpf.

Where applicable, adult experiments used either mixed-sex cohorts or sex-stratified groups as indicated in the figure legends. Larval experiments were performed at stages prior to sexual differentiation. Experiments were performed in both larval and adult zebrafish to match assay requirements to biological context. Larval models were used where optical transparency, whole-mount imaging, germ-free derivation, and short-term pharmacologic manipulation are required. Adult models were used to assess mature intestinal architecture, long-term epithelial homeostasis, microbiome composition, and sex-dependent phenotypes. Developmental stage is specified for each experiment in the figure legends and indicated by icons in figure panels.

### Generation and validation of the *nod2*-mutant zebrafish line

crRNA:tracrRNA duplex and gRNA:Cas9 RNP complexes were prepared with the *nod2* crRNA sequence: TTGGACCTGCTACTTGCTC. Generation and validation of the *nod2*-mutant zebrafish line was performed as previously described [31], with modifications detailed below. Briefly, Wild-type TL zebrafish embryos were microinjected at the 1-4-cell stage with 1 nL of the CRISPR-Cas9 injection mix per embryo, using a reduced injection volume compared to the original protocol. At 5 days post-fertilization (dpf), larvae were collected for genomic DNA extraction. The genomic region encompassing the CRISPR target site was amplified by PCR, purified, and subjected to Sanger sequencing to assess mutation efficiency. Embryos with evidence of CRISPR-Cas9-induced mutations were raised to adulthood and outcrossed to wild-type TL zebrafish. Genomic DNA was extracted from F1 larvae at 5 dpf and screened for germline transmission of *nod2* mutations using high-resolution melting (HRM) analysis. Clutches positive for mutations were raised to adulthood, and heterozygous carriers were identified by HRM, PCR, and Sanger sequencing. Founder fish harboring the same mutation were selected and in-crossed, and F2 embryos were sequenced to confirm the presence of the intended mutation. The resultant mutant line carried a 4-base pair deletion (CTTG) in exon 2 of *nod2*, resulting in a frameshift and truncated, non-functional protein.

### Fish weight, body length, and gut length measurement

Adult zebrafish were anesthetized in Tricaine (MS-222) until complete loss of touch response and then blotted gently on absorbent paper to remove excess water. Body weight was recorded immediately using a precision analytical balance (+/- 0.1 mg sensitivity), with each individual weighed prior to any dissection to avoid dehydration artifacts. Standard body length was measured from the tip of the snout to the base of the caudal fin from lateral images acquired under a dissection microscope using an adjacent metric ruler for calibration.

For gut length measurements, fish were euthanized, and the gastrointestinal tract was carefully dissected under a stereomicroscope using fine forceps. Surrounding tissue was removed without stretching the tissue. The intestine was then laid flat in phosphate-buffered saline (PBS) in a glass Petri dish and imaged with a ruler under a dissection scope. Total gut length was quantified using ImageJ, measuring the longest continuous distance from anterior to distal end. All measurements were performed blinded to genotype, and at least three technical measurements per intestine were averaged to account for handling variability. For size-normalized analyses, gut length was divided by body length or body weight, as indicated in the figure panels.

### Generating single-cell suspensions for single cell RNA-sequencing

Single-cell suspensions were prepared from adult zebrafish intestines as previously described [23], with minor modifications. Briefly, five adult WT TL and five adult *nod2^-/-^* cohoused in-crossed siblings with equal sex distribution (2 males:3 females) were selected for tissue collection. Intestines were dissected, minced, and digested in a dissociation cocktail containing fresh collagenase A (1 mg/mL), 40μg/mL Proteinase K, and 10X Trypsin (0.25% final). Following digestion, cell suspensions were washed, filtered, and live cells isolated using OptiPrep Density Gradient Medium according to OptiPrep Application Sheet C13. Viable cells were counted and processed for single-cell RNA sequencing as described [23]. Cell suspensions were then diluted to a concentration of 1200 cell/μL and immediately run through the 10X Genomics Chromium Controller with Chromium Single Cell 3′ Library & Gel Bead Kit v3.1. Library QC and sequencing was performed by Novogene using the Illumina HiSeq platform.

### Processing and analysis of single-cell RNA-seq data

Single-cell RNA-seq data were processed and analyzed as previously described [23], with minor modifications. Briefly, Cell Ranger v7.0.1 (10X Genomics) was used to demultiplex raw sequencing files and align reads to the zebrafish reference genome (Ensembl GRCz11.96). Cell Ranger output matrices were analyzed using the Seurat R package (v5.0.1) in RStudio. Cells with fewer than 200 unique molecular identifiers (UMIs), greater than 2500 UMIs, or greater than 25% mitochondrial reads were removed. Seurat was used to normalize expression values and perform clustering (resolution 0.8, 50 principal components), with optimal PCs determined by JackStraw scores and elbow plots. Marker genes were identified using the “FindMarkers” function, and clusters were annotated using known zebrafish cell type markers or mammalian orthologs.

### 16S rRNA gene sequencing

Briefly, four adult WT TL and four *nod2^-/-^* 6-month-old in-crossed siblings with equal sex distribution were selected for microbiome analysis. For the co-housing condition, WT and *nod2^-/-^* adults were maintained together in the same recirculating tank for ≥8 weeks prior to sampling to ensure shared environmental microbiota. For the single-housing condition, WT and *nod2^-/-^* adults were maintained in separate tanks on the same recirculating water system under identical water quality, feeding schedule, and density for the same duration; despite shared system water, tank-level separation can still permit divergence in gut microbial communities. All fish were fasted prior to dissections. Zebrafish intestines were dissected with surrounding organs removed. Individual guts were collected and homogenized in 300μL of Powerbead Solution. The Qiagen DNeasy UltraClean Microbial Kit (Cat No. 12224-250) was used to extract microbial DNA. To assess the intestinal bacterial community composition, the V4 variable region of the 16s rRNA gene encompassed by the 515 forward primer and 806 reverse primer was sequenced. Quality control and sequencing was performed by Novogene Corporation using illumina Novaseq Platform PE250. Sequences were processed with QIIME2-2024.10. The DADA2 pipeline was used to join paired-end reads, remove chimeric sequences, and to generate the feature table used to resolve amplicon sequence variants using default parameters. DADA2 denoising resulted in 694,045 reads for co-housed and 814,688 reads for single-housed. Amplicon sequence variants (ASVs) represented by fewer than 200 reads across all samples were removed. A naïve Bayes classifier trained on SILVA138 99% full-length reference sequences was used to assign taxonomy. Taxonomy assignments were verified using NCBI blast. The sequence table was then filtered to exclude any sequences that were unassigned, not assigned past phylum level, or assigned as eukaryota, resulting in 575,576 sequences (co-housed) or 673,162 sequences (single-housed) corresponding to 25 unique features.

### RNA extraction and cDNA Synthesis

Total RNA was extracted from zebrafish intestinal tissue using TRIzol Reagent (Thermo Fisher Scientific) following manufacturer protocols with minor modifications. Intestinal samples were collected from five WT TL males, five WT TL females, five *nod2*^-/-^ males, and five *nod2*^-/-^ females, all derived from cohoused in-crossed sibling groups. Fish were fasted prior to dissection. Individual intestines were isolated, cleared of surrounding tissue, and homogenized in 250 μL of TRIzol using a motorized pestle. Homogenates were brought to 1 mL total volume with additional TRIzol and stored at -70 °C until further processing. For phase separation, 400 μL of chloroform was added, followed by 15 s vortexing and a 3-minute incubation at room temperature. The aqueous phase was collected, and chloroform extraction was repeated at a 1:1 ratio. RNA was precipitated by adding 50 μL of 3 M sodium acetate (pH 5.2) and 1 mL of 95% ethanol, followed by overnight incubation at 4 °C. RNA was pelleted by centrifugation (12,000 × g, 10 min, 4 °C), washed twice in 0.5 mL 75% ethanol (7,500 xg, 10 min, 4 °C), and resuspended in 50 μL of nuclease-free water. RNA concentrations were determined using a NanoDrop spectrophotometer and diluted to a standardized concentration for downstream cDNA synthesis. cDNA was synthesized from up to 3 μg of total RNA using the qScript cDNA SuperMix (Quantabio, no. 95048-100) in a total reaction volume of 20 μL. Reactions were incubated at 25 °C for 5 min, 42 °C for 30 min, and 85 °C for 5 min, followed by a 4 °C hold.

### Quantitative RT-qPCR

Quantitative PCR was performed using PerfeCTa SYBR Green SuperMix (Quantabio, no. 95054-500) on a One-Step real-time PCR thermocycler. Reactions were carried out in a 10 μL volume containing 2.5 μL of cDNA and 1.6 μM of each forward and reverse primer. Plates were centrifuged at 200 × g for 1 minute prior to thermocycling. The qPCR program consisted of an initial denaturation at 95 °C for 3 minutes, followed by 40 amplification cycles of 95 °C for 15 seconds and 60 °C for 45 seconds, with melt curve analysis included at the end of the run. All reactions were performed in technical triplicates. Cycle threshold (Ct) values were averaged across replicates, and relative gene expression was calculated using the ΔΔCt method, with *ef1α* used as the endogenous reference gene.

### Zebrafish RT-qPCR primer sequences

The following sequences were used (given in order from the 5′ to 3′ end): *vtg1* qPCR forward, GCCAAAAAGCTGGGTAAACA; *vtg1* qPCR reverse, AGTTCCGTCTGGATTGATGG; *vtg2* qPCR forward, TGCCGCATGAAACTTGAATCT; *vtg2* qPCR reverse, GTTCTTACTGGTGCACAGCC; *vtg4* qPCR forward, TCCAGACGGTACTTTCACCA; *vtg4* qPCR reverse, CTGACAGTTCTGCATCAACACA; *ef1a* qPCR forward, GCATACACTAAGAAGATCGGC; *ef1a* qPCR reverse, TCTTCCATCCCTTGAACCAG.

### Generating germ-free zebrafish

Fish embryos were made germ-free as described in [52]. A clutch of embryos was collected then washed and split into two cohorts. The CV cohort was kept in sterile EM, while the GF cohort was kept in sterile EM supplemented with ampicillin (100 μg/mL), kanamycin (5 μg/mL), amphotericin B (250 ng/mL), and gentamicin (50 μg/mL). Embryos were washed every 2 hours with EM or EM plus antibiotics for CV and GF cohorts respectively. Once at 50% epiboly, the GF cohort was successively washed three times in EM, then 2 minutes in 0.1% polyvinylpyrrolidone-iodine (PVP-I) in EM, followed by three EM washes, then a 20-minute incubation with 0.003% sodium hypochlorite (bleach) in EM. Embryos were washed three more times then transferred into tissue culture flasks with sterile EM. The CV cohort received the same number and duration of washes, using EM in lieu of dilute PVP-I or bleach. All work was performed in a biosafety cabinet sterilized first with 10% bleach, followed by 70% ethanol. We tested for bacterial contamination in GF flasks at 4 days post-fertilization, according to established protocol. EM was collected from CV and GF culture flasks to test for bacteria by plating on TSA. Parental tank water and sterile filtered water were used as a positive and negative control respectively, where bacteria were positively identified in parental tank water and confirmed absent from sterile water. CV and GF flasks with bacteria present or absent respectively were used for subsequent analysis.

### Flow Cytometry of intestinal proliferation in zebrafish larvae

To quantify intestinal epithelial proliferation, zebrafish larvae were immersed in 100 μg/mL 5-ethynyl-2’-deoxyuridine (EdU; 10 mM stock in DMSO) for 24 hours in embryo medium containing 1% DMSO, with or without muramyl dipeptide (MDP) at 1 μg/mL. A 10 mg/ml working stock of MDP (Sigma-Aldrich A9519) was diluted to 1 μg/ml for all treatments. For MDP treatments, axenic larvae were exposed to MDP for 16 hrs prior to EdU labeling. Following treatment, intestines were dissected from 15 larvae per condition in ice-cold PBS and subjected to enzymatic dissociation in a cocktail containing 1 mg/mL collagenase, 0.25% trypsin, and 40 μg/mL proteinase K at 37°C for 30 minutes with intermittent gentle trituration. Dissociated cells were filtered through a 40 μm strainer, pelleted by centrifugation (300 × g, 10 min, 4°C), and washed with PBS. Cells were fixed in 4% paraformaldehyde for 15 minutes at room temperature, washed with FACS buffer (PBS without calcium or magnesium supplemented with 2% heat-inactivated calf serum), and permeabilized in saponin buffer (0.1% saponin in FACS buffer) for 10 minutes. EdU detection was performed using the Click-iT® EdU Alexa Fluor 488 Imaging Kit according to the manufacturer’s instructions. After EdU labeling, cells were stained with DRAQ5 (1:1000) for nuclear visualization. Flow cytometry was conducted on the Attune NxT cytometer, and EdU⁺ proliferating cells were quantified based on fluorescence intensity using appropriate controls for gating.

### Estrogen and Tamoxifen treatment of zebrafish larvae

A working stock of 20mg/mL 17β-estradiol (E2; Sigma) was prepared in DMSO and diluted to a final concentration of 25 ng/µL in 1xEM for all treatments. No more than 20 zebrafish larvae at 5 dpf were placed in each petri dish and incubated in 25 ng/µL E2 or control DMSO for 24 hrs. Larvae were then collected for downstream applications, including immunofluorescence and whole-mount Alcian blue staining. Similarly, a 20mg/mL stock of tamoxifen (Sigma), dissolved in DMSO, was diluted to a final concentration of 1.5 ng/µL in 1xEM. Larvae (5 dpf; ≤20 per dish) were incubated in 1.5 ng/µL tamoxifen for 6 hrs prior to collection for EdU labeling and Alcian blue staining. For DSS survival assays, larvae were pretreated with E2 for 24 hrs or tamoxifen for 6 hrs before initiating DSS exposure.

### DSS treatment of zebrafish larvae

DSS treatments were performed as previously described [32], with 0.5% DSS (Sigma, cat. no. 42867) used for all experiments and treatments initiated at 5 dpf. Briefly, no more than 20 zebrafish larvae were placed in each petri dish and incubated in freshly prepared 0.5% DSS dissolved in 1x EM for 24 hrs. Larvae were collected for further experimental use after removal of DSS. For survival assays, larvae were continuously exposed to 0.5% DSS in 1x EM beginning at 5 dpf. Media was replaced daily by transferring larvae into fresh DSS-containing petri dishes. Survival was monitored over time and quantified by Kaplan-Meier analysis, using 20 larvae per genotype per condition

### Whole-mount Alcian blue staining and quantification

This protocol was adapted and modified from [32]. To visualize goblet cells, zebrafish larvae (7 dpf) were fixed in 4% PFA at 4 °C for 48 hours, rinsed in 1x PBS for 5 min, and incubated in acidic ethanol (70% ethanol containing 1% concentrated HCl) for 10 min. Alcian blue stains secreted acidic mucins rather than goblet cells directly; therefore, this assay quantifies mucin-positive area and does not measure goblet cell number or density. Larvae were stained in 0.1% Alcian blue (prepared in 3% acetic acid, pH 2.5) for 3 hours at room temperature, washed three times in acidic ethanol, and incubated overnight in acidic ethanol to reduce background staining. Larvae were mounted in 0.7% low-melting-point agarose and imaged under identical illumination and exposure settings. For quantification, images were analyzed in Fiji (ImageJ) using blinded analysis. A region of interest (ROI) corresponding to the mid and distal intestinal segments, defined anteriorly by the hepatic bend and posteriorly by the cloaca, was outlined while excluding extra-intestinal tissues. Images were converted to RGB, and a single fixed color threshold for Alcian blue signal was established using control images and then applied uniformly to all samples within the experiment without per-image adjustment. Threshold values were held constant across genotypes and treatments to avoid segmentation bias. Alcian blue-positive regions were segmented using this fixed threshold, and the Alcian blue-positive area and total ROI area were measured using the Analyze Particles tool. Data are reported as percent Alcian blue-positive intestinal area, calculated as (AB^+^ area/total intestinal ROI area) X 100, reflecting the proportion of the intestinal surface occupied by goblet cell-associated mucin staining. Each dot in quantification plots represents a single larval intestine.

### Immunofluorescence and fluorescence microscopy

Adult zebrafish intestines were fixed in 4% PFA (diluted in 1xPBS) overnight at 4°C. Intestines were washed twice in 1xPBS. Intestinal segments were then cryoprotected in 15% (w/v) sucrose/PBS at room temperature until sunk followed by sinking in 30% (w/v) sucrose/PBS (overnight at 4°C). Intestines were mounted in Optimal Cutting Temperature Embedding Medium, then frozen on dry ice. Seven-micron cryosections were collected on Superfrost Plus slides. After allowing sections to adhere onto slides, slides were immersed in PBS for 20 min at room temperature. Tissue was blocked for 1hr at room temperature in 3% (w/v) BSA dissolved in PBST (1xPBS +0.2% (v/v) Tween 20). Primary antibodies were diluted in blocking buffer. Sections were incubated in primary antibody solution overnight in a humid chamber at 4°C. The primary antibodies used in this paper was mouse anti-zebrafish gut secretory cell epitopes antibody [FIS 2F11/2] 1/500 (abcam 71286), and chicken polyclonal anti-GFP (ThermoFisher Scientific Cat# PA1-9533). Excess primary antibody was washed by immersing slides in fresh PBST for 1.5hrs. Sections were incubated in secondary antibody solution (prepared in blocking buffer) for 1hr at room temperature in a humid chamber, protected from light. To stain apoptotic cells, secondary solution was removed then sections incubated in TUNEL reaction mixture following the manufacturer’s suggestion (Roche 12156792910). Lastly, nuclei were stained with Hoechst (Molecular Probes Cat# H-3569) diluted 1:2000 in PBST for 10 min protected from light. Slides were washed in PBST by brief immersion followed by a 30-min incubation in fresh PBST protected from light. Slides were mounted in Fluoromount Aqueous Mounting Medium.

Zebrafish larvae (6-7 dpf) were incubated in embryo medium with 1% DMSO and 5 mM EdU for 8 hrs at 29°C. Fish intestines were dissected in PBS and fixed in 4% paraformaldehyde in PBS overnight at 4°C. Guts were washed three times with PBSTx (0.75% Triton X-100 in 1X PBS), blocked for 1 hr in PBSTx +3% BSA at room temperature, and stained with primary antibodies in blocking buffer overnight at 4°C. Guts were washed with PBSTx and then stained for 1 hr at room temperature with secondary antibodies and nuclear stain (1/1000 Hoechst 33258), followed by rinse in PBSTx. EdU detection was performed by incubating guts in Click-iT reaction cocktail for 30 min at room temperature. Guts were washed in PBSTx followed by extra washing in PBS. Whole larval intestines were mounted on microscope slides using Flouromount and visualized.

### EdU proliferation quantification

For imaging-based proliferation assays, whole-mount intestines were imaged by confocal microscopy using identical acquisition settings across experimental groups. EdU⁺ nuclei were quantified either across the entire intestine (baseline genotype comparisons) or within a standardized intestinal region of interest (ROI) spanning the mid-intestine (rearing-condition comparisons), depending on experimental design. Nuclei were identified by DNA staining and EdU⁺ cells were counted using blinded analysis. Percent EdU⁺ values were calculated relative to total epithelial nuclei within the quantified region. Flow cytometry-based EdU quantification was performed on dissociated intestines where indicated to enable unbiased enumeration following acute ligand stimulation.

### Quantification of Anxa4⁺, Lcp1⁺, and TUNEL⁺ cells

Quantification of Anxa4⁺ secretory cells, Lcp1⁺ leukocytes, and TUNEL⁺ apoptotic cells was performed on sagittal intestinal sections using Fiji (ImageJ). Individual villi were manually outlined using the DNA (nuclear) channel to define the epithelial region of interest. Within each villus ROI, Anxa4⁺, Lcp1⁺, and TUNEL⁺ cells were detected using a fixed threshold applied uniformly across all samples and counted manually using the cell counter tool. Total epithelial cell number per villus was determined by counting all DNA⁺ nuclei within the same ROI. To account for differences in villus size and cellular density between sections and fish, data were expressed as the percentage of marker-positive cells per villus, calculated as (number of Anxa4⁺, Lcp1⁺, or TUNEL⁺ cells ÷ total number of DNA⁺ nuclei within the ROI) x 100.

### Immunohistochemistry

Adult zebrafish intestines were fixed at 4°C in BT fixative: 4% PFA, 0.15mM CaCl2, 4% Sucrose in 0.1 M phosphate buffer (pH 7.4). Intestinal segments were processed for paraffin embedding and 5μm sections collected on Superfrost Plus slides. Tissue was deparaffinized, rehydrated, then boiled in 10mM sodium citrate pH 6, 0.05% Tween 20 for 20 min at 98°C to unmask antigen. After cooling sections to room temperature for 30 min, sections were incubated in 3% hydrogen peroxide for 10 min, washed twice in dH2O, and once in PBSt (PBS +0.5% Triton X-100). Sections were then blocked for 1hr at room temperature in 10% (v/v) normal goat serum (NGS)/PBSt followed by overnight incubation in primary antibody solution (prepared in blocking buffer) in a humid chamber at 4°C. The primary antibodies used in this paper were: mouse anti-PCNA 1:5000 (Sigma P8825), rabbit anti-Lcp1 1:1000 (GeneTex GTX124420) and a custom anti-zebrafish NOD2 antibody (1:1000, Biologics International Corp (BIC)). Slides were washed three times in PBSt and tissue incubated in SignalStain Boost Detection Reagent (HRP, Rabbit or Mouse) for 30 min at room temperature. Colorimetric detection was done by incubating in SignalStain DAB Chromogen solution for 5 min. Sections were counterstained with Alcian blue (Aldrich 19,978-8) for 3 mins, counterstained in quarter-strength Hematoxylin Gill III (Leica Ca.# 3801542) for 30 s, dehydrated in ethanol, cleared with toluene and mounted in DPX.

### Microscopy

Histological samples were imaged on a ZEISS AXIO A1 compound light microscope with a SeBaCam 5.1MP camera. Confocal images were captured on an Olympus IX-81 microscope fitted with a CSU-X1 spinning disk confocal scan-head, Hamamatsu EMCCD (C9100-13) camera and operated with PerkinElmer’s Volocity. Confocal z stack image processing was done in Fiji.

### Quantification and statistical analysis

Statistical analyses were performed using R. The statistical tests, group statistics, and P values used in each experiment are described in the respective figure legends.

### Data and code availability

- Single-cell RNA-seq raw data files have been deposited to the NCBI GEO database and are publicly available (NCBI GEO (GSE299263)).
- This paper does not report original code.
- Any additional information required to reanalyze the data reported in this work paper is available from the lead contact upon request.

## ACKNOWLEDGMENTS

We acknowledge support with single-cell library preparation from Mike Wong at the Faculty of Medicine and Dentistry Advanced Cell Exploration (ACE) Core. We acknowledge flow cytometry support from Dr. Aja Rieger, Rikus Niemand, Lai Xu, and Xinyue Xu. Flow Cytometry Facility experiments were performed at the University of Alberta Faculty of Medicine & Dentistry Flow Cytometry Facility, which receives financial support from the Faculty of Medicine & Dentistry and Canada Foundation for Innovation (CFI) awards to contributing investigators. We acknowledge microscopy support from Dr. Hilmar Strickfaden, and Kiera Smith at the Faculty of Medicine and Dentistry Cell Imaging Core. We acknowledge the immunohistochemistry support provided by Dr. Kacie Norton at the Department of Biological Sciences Advanced Microscopy Facility. We would also like to thank Science Animal Support Services and the Alberta Health Sciences Animal Laboratory Services for their excellent care of the zebrafish aquatic facilities. We acknowledge Sanger DNA sequencing and HRM analysis support from Troy Locke at the Facility of Science Molecular Biology Facility (MBSU). Thanks to Dr. Dana Philpott, Derek Tsang, Dr. Lisa Willis, Lena Jones, Dr. Aurélia Joly, and Dr. Reegan Willms for critical reading of the manuscript. This work was supported by a grant from the Canadian Institutes of Health Research (MOP77746) to E.F. M.E. has funding support through the Boytzun Studentship in Colorectal Cancer, Cancer Research Institute of Northern Alberta (CRINA), and Li Ka Shing Institute of Virology Graduate Studies Entrance Award.

## AUTHOR CONTRIBUTIONS

M.E. and E.F. conceived and designed the experiments. M.E carried out the experiments. M.E. and E.F. performed computational analysis. Funding for this project was acquired by E.F. M.E. and E.F. wrote the manuscript.

## DECLARATION OF INTERESTS

The authors declare no competing interests.

## Notes

### Competing Interest Statement

The authors have declared no competing interest.

### Summary of Updates

- Clarification of various methods. - Additional data stratifying phenotypes by sex. - Expanded discussion of limitations and future directions.

